# Swirling motion of breast cancer cells radially aligns collagen fibers to enable collective invasion

**DOI:** 10.1101/2025.01.31.635980

**Authors:** Aashrith Saraswathibhatla, Md Foysal Rabbi, Sushama Varma, Vasudha Srivastava, Olga Ilina, Naomi Hassan Kahtan Alyafei, Louis Hodgson, Zev Gartner, Peter Friedl, Robert West, Taeyoon Kim, Ovijit Chaudhuri

## Abstract

In breast cancer (BC), radial alignment of collagen fibers at the tumor-matrix interface facilitates collective invasion of cancer cells into the surrounding stromal matrix, a critical step toward metastasis. Collagen remodeling is driven by proteases and cellular forces, mediated by matrix mechanical plasticity, or irreversible matrix deformation in response to force. However, the specific mechanisms causing collagen radial alignment remain unclear. Here, we study collective invasion of BC tumor spheroids in collagen-rich matrices. Increasing plasticity to BC-relevant ranges facilitates invasion, with increasing stiffness potentiating a transition from single cell to collective invasion. At enhanced plasticity, cells radially align collagen at the tumor-matrix interface prior to invasion. Surprisingly, cells migrate tangentially to the tumor-matrix interface in a swirling-like motion, perpendicular to the direction of alignment. Mechanistically, swirling generates local shear stresses, leading to distally propagating contractile radial stresses due to negative normal stress, an underappreciated property of collagen-rich matrices. These contractile stresses align collagen fibers radially, facilitating collective invasion. The basement membrane (BM), which separates epithelia from stroma in healthy tissues, acts as a mechanical insulator by preventing swirling cells from aligning collagen. Thus, after breaching the BM, swirling of BC cells at the tumor-stroma interface radially aligns collagen to facilitate invasion.

## Introduction

Collective invasion of BC cells from a pre-invasive carcinoma in situ into the surrounding collagen (col1)-rich stromal matrix represents a critical step towards metastasis, with changes in the architecture and mechanics of the col1-rich stromal matrix implicated as playing a key role. Increased stiffness of the col1-matrix occurring during invasive BC (IBC) progression promotes an invasive phenotype in the carcinoma cells, and denser col1 promotes collective invasion over single-cell invasion^1–6^. Further, it has long been known that the stromal col1 fibers become aligned radially with respect to the tumor-stroma interface, an architecture defined as the tumor-associated col1 structure-3 (TACS-3)^5,7–10^ (Fig. 1a-b). Radial alignment of col1 has been implicated in guiding the collective invasion of cancer cells out of the tumor by providing tracks for invasion. Physiologically, TACS-3 has prognostic significance for metastasis^10^, patient survival^8^, and BC relapse post-mastectomy^11^, marking it as a critical event in BC pathogenesis.

**Figure 1:**
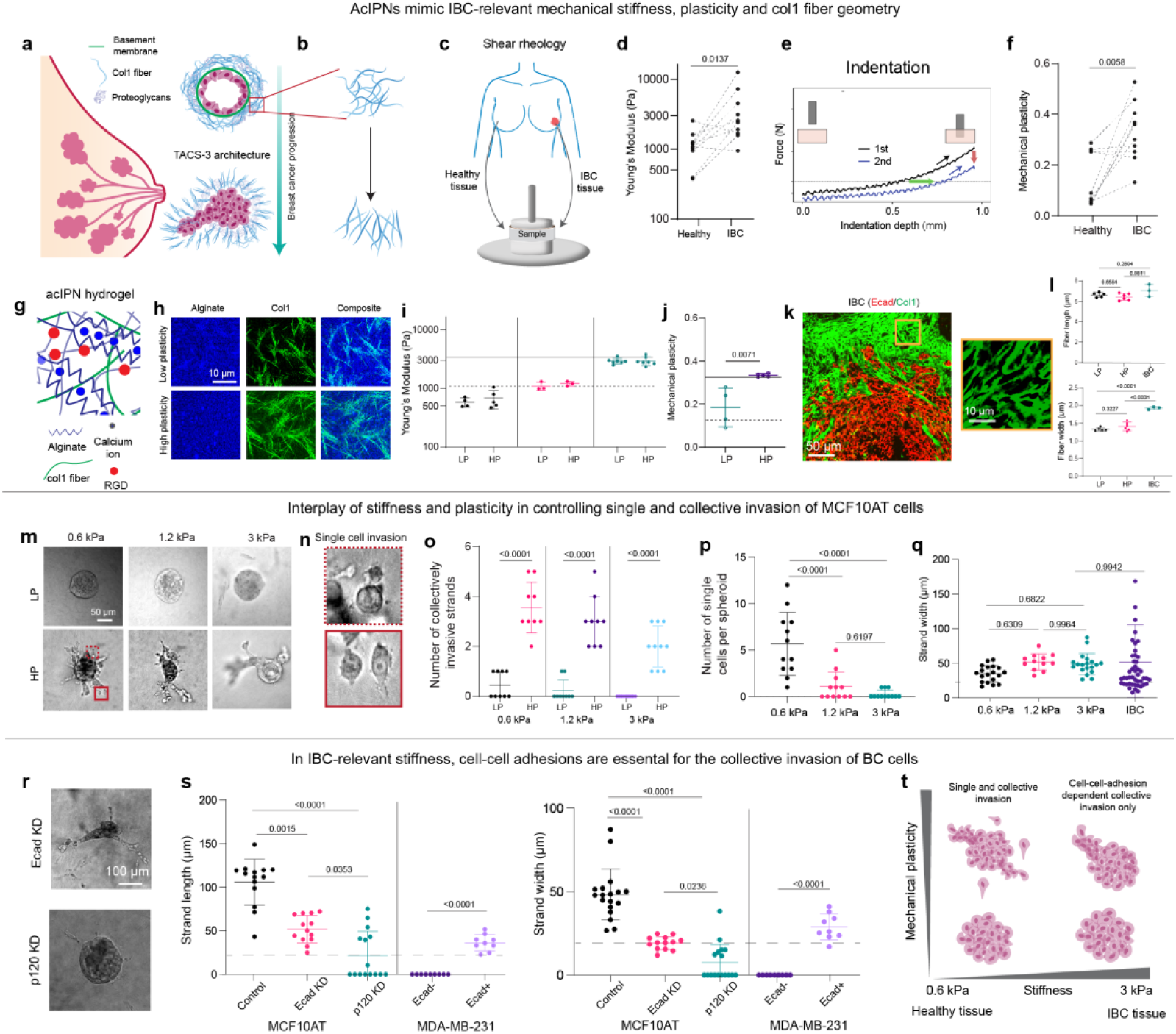
In col1 rich ECMs, enhanced plasticity increases invasion of BC cells, with enhanced stiffness promoting collective over single cell invasion. **a**, Schematic of the progression of BC, where a collective group of cells invade along the aligned col1 at tumor-stroma interface after breaching the basement membrane. **b**, Schematic of col1 fibers showing random and aligned configurations. **c**, Schematic of rheological measurements of human healthy and IBC tissues. **d**, Young’s modulus of healthy and their counterpart IBC tissues (n=9 patients). **e**, Schematic of force vs. indentation depth curves of two different cycles separated by one hour. The green arrow indicates permanent deformation, and red arrow indicates the drop in peak force. Dotted line corresponds to 25% of peak force in the first indentation curve. Figure adapted from Ref. 16. **f**, Mechanical plasticity of healthy and their counterpart IBC tissues (n=9 patients). **g**, Schematic of an interpenetrating network of type-1 collagen and alginate (acIPN). **h**, Airy Scan imaging of fluorescently tagged alginate and col1, and their overlay. **i**, Young’s modulus of LP and HP acIPNs of three different acIPNs (n≥3 different hydrogels). Horizontal dotted and solid lines correspond to average Young’s modulus of healthy and IBC tissues respectively. **j**, Mechanical plasticity of LP and HP acIPNs at 3 kPa stiffness (n≥3 hydrogels). Horizontal dotted and solid lines correspond to average plasticity of healthy and IBC tissues respectively. **k**, (Left) Fluorescent staining of Ecad and col1 in an IBC histopathology section. (Right) Zoomed-in image of the col1 fiber architecture in the section. **l**, Average length and width of col1 fibers in LP and HP acIPNS and in IBC tissue sections (n≥3 hydrogels or 3 patients). **m**, Phase contrast images of MCF10AT spheroids on day 5 in LP and HP acIPNs of 0.6 kPa, 1.2 kPa and 3 kPa stiffness. **n**, Closeup of single cell invasions in HP acIPN of 0.6 kPa stiffness. **o**, Number of collectively invading strands from MCF10AT spheroids on day 5 in LP and HP acIPNs of 0.6 kPa, 1.2 kPa and 3 kPa (n≥ 9 spheroids; N=3 biological replicates). **p**, Number of single cells in HP acIPNs of three different stiffness conditions (n≥ 9 spheroids; N = 3). **q**, Average width of the collectively invading strands in HP acIPNs of 3 different stiffness conditions and in IBC condition (n≥ 9 spheroids; N = 3). Data for IBC was taken from Ref. 6. The dotted line indicates the average length of a single cell. **r**, Phase contrast images of Ecad KD and p120 KD spheroids on day 5 in HP acIPN of 3 kPa stiffness. **s**, Average length and width of collectively invading strands in control, Ecad KD and p120 KD conditions of MCF10AT spheroids, and in Ecad- and Ecad+ conditions of MDA-MB-231 spheroids (n≥ 9 spheroids; N=3). The dotted line indicates the average length of a single cell (n≥ 9 spheroids; N = 3). **t**, A schematic summarizing the roles of stiffness, plasticity and cell-cell adhesions in controlling the invasion of BC cells. Panels **d -** Wilcoxon matched pair signed rank test. Panel **f** – Two-sided t-test. Panels **l, p, q s -** One-way variance (ANOVA) with Tukey’s correction for multiple groups. Panel **o, s** – Two-sided t-test for comparing two groups.

Previous studies have shown that cancer-associated fibroblasts (CAFs) can remodel col1 at the tumor-matrix interface into a TACS-3-like architecture, using proteases and contractile mechanical forces to guide the invasion of BC cells^6,9,12,13^. However, it is also possible that the BC cells can autonomously implement TACS-3-like remodeling in collaboration with or independently of the CAFs. Indeed, cancer cells are known to secrete proteases, particularly matrix metalloproteinases (MMPs), to degrade and remodel the matrix during collective invasion^14^. In addition, cancer cells can physically remodel matrices through mechanical forces if that matrix exhibits sufficient matrix mechanical plasticity, or irreversible deformations in response to mechanical forces^15–17^. Here, we investigate the role of matrix stiffness and mechanical plasticity of col1-rich ECMs in controlling the collective invasion of BC cells and whether and how cancer cells collectively remodel the col1-rich ECM into TACS-3-like configuration to facilitate collective invasion.

## Results

### Alginate-col1 interpenetrating network hydrogels mimic the stromal matrix in BC tissues

We first assessed the mechanical properties of healthy tissues, and their counterpart IBC tissues (Fig. 1c). In rheological measurements of stiffness, IBC tissues showed a significant increase in average stiffness (∼4.7 kPa) compared to healthy tissues (∼1.4 kPa) (Fig. 1d), consistent with the findings of previous studies^3,5,18^. Next, tissue mechanical plasticity was measured using mechanical indentation and assessing the extent to which the indentation induced permanent deformation in the tissue (Fig. 1e)^16^. IBC tissues exhibited a significant increase in mechanical plasticity (32%) compared to their healthy tissues (15%) (Fig. 1f). Overall, these measurements establish the relevance of both increased stiffness and matrix mechanical plasticity during BC progression.

Guided by these tissue-scale measurements, we designed col1-rich hydrogels to mimic the stiffness and mechanical plasticity of healthy and IBC tissues tissue (Fig. 1d, f). The stromal ECM consists of both a fibrous col1 network as well as other interstitial components such as hyaluronic acid and fibronectin, making it nanoporous. Therefore, interpenetrating networks of alginate and col1 (acIPNs) were used to mimic both the fibrous col1 architecture and overall nanoporosity of stromal ECM while providing mechanical tunability (Fig. 1g-h). Alginate is a polymer derived from seaweed that is inert to any cell adhesion receptors, is not susceptible to cell-secreted proteases, and can be cross-linked ionically with divalent cations (calcium) into a nanoporous hydrogel4. In the acIPNs utilized here, alginate is conjugated with the cell adhesion peptide motif RGD (arginine– glycine–aspartic acid) to facilitate adhesion to the alginate, and the molecular weight (MW) and cross-linker concentration of alginate are varied in order to modulate stiffness and plasticity independently^15,16^.

AcIPNs were formed with three different stiffnesses (0.6, 1.2 and 3 kPa), with 0.6 kPa and 3 kPa capturing the stiffness of healthy and IBC tissues, respectively (Fig. 1i). At each stiffness, acIPNs were formed that exhibited either low plasticity (LP) or high plasticity (HP), matching that of healthy and IBC tissues respectively (Fig. 1j; Extended Fig. 1a). For all acIPNs, the col1 concentration was kept at 1.5 mg/mL, and col1 fiber length was comparable to an invasive BC (IBC) tissue, though the col1 fiber thickness was slightly higher in BC tissues (Fig. 1k-l; Extended Fig. 1b-e). Thus, the stiffness and plasticity of acIPNs can be tuned independently over the range relevant to healthy and IBC tissues, while partially capturing the col1 fiber geometry of IBC tissues.

### Collective invasion requires sufficient matrix mechanical plasticity and is promoted by increased stiffness

Next, we investigated how the stiffness and plasticity of acIPNs regulated the invasion of BC spheroids. Spheroids of MCF10AT cells, an invasive cell line whose invasion is collective, were encapsulated in the LP and HP acIPNs in all three stiffness conditions, and invasion was assessed on day 5. Across the entire range of stiffness, spheroids invaded robustly in HP acIPNs, with both single-cell invasions and strands of collective cells invading the HP acIPNs but did not invade in LP acIPNs (Fig. 1m-p; Extended Fig. 2a; Supplementary Video 1). These indicate that sufficient matrix mechanical plasticity is required for BC invasion independent of stiffness. Interestingly, 0.6 kPa HP acIPNs exhibited a significant number of single-cell invasions with hardly any single-cell invasions in 3 kPa HP acIPN. This is similar to recent findings that increased confinement due to higher col1 density promoted collective over single-cell invasion, though col1 was kept constant while changing the stiffness in our assays^6,19^ (Fig. 1n, p). The dominant mode of invasion observed was strands of collectively invading cells with lengths and widths more than the average length of a single cell (Fig. 1o, q; Extended Fig. 2b). Importantly, the strand widths were of similar length scales to the widths of collectively invading strands previously reported in human IBC (Fig. 1q)^6^. Similar results on the impact of plasticity were observed in 4T1 cell spheroids were observed (Extended Fig. 2c-d). Further, MCF10AT spheroids expressed E-cadherin (Ecad), indicating that they maintain their cell-cell junctions. However, they also display vimentin, indicating that cells maintained their cell-cell junctions but may display some mesenchymal characteristics (Extended Fig. 2e-f). Altogether, we demonstrate that matrix mechanical plasticity is required for the invasion of BC cells and that increased stiffness promotes collective invasion over single-cell invasion.

**Figure 2:**
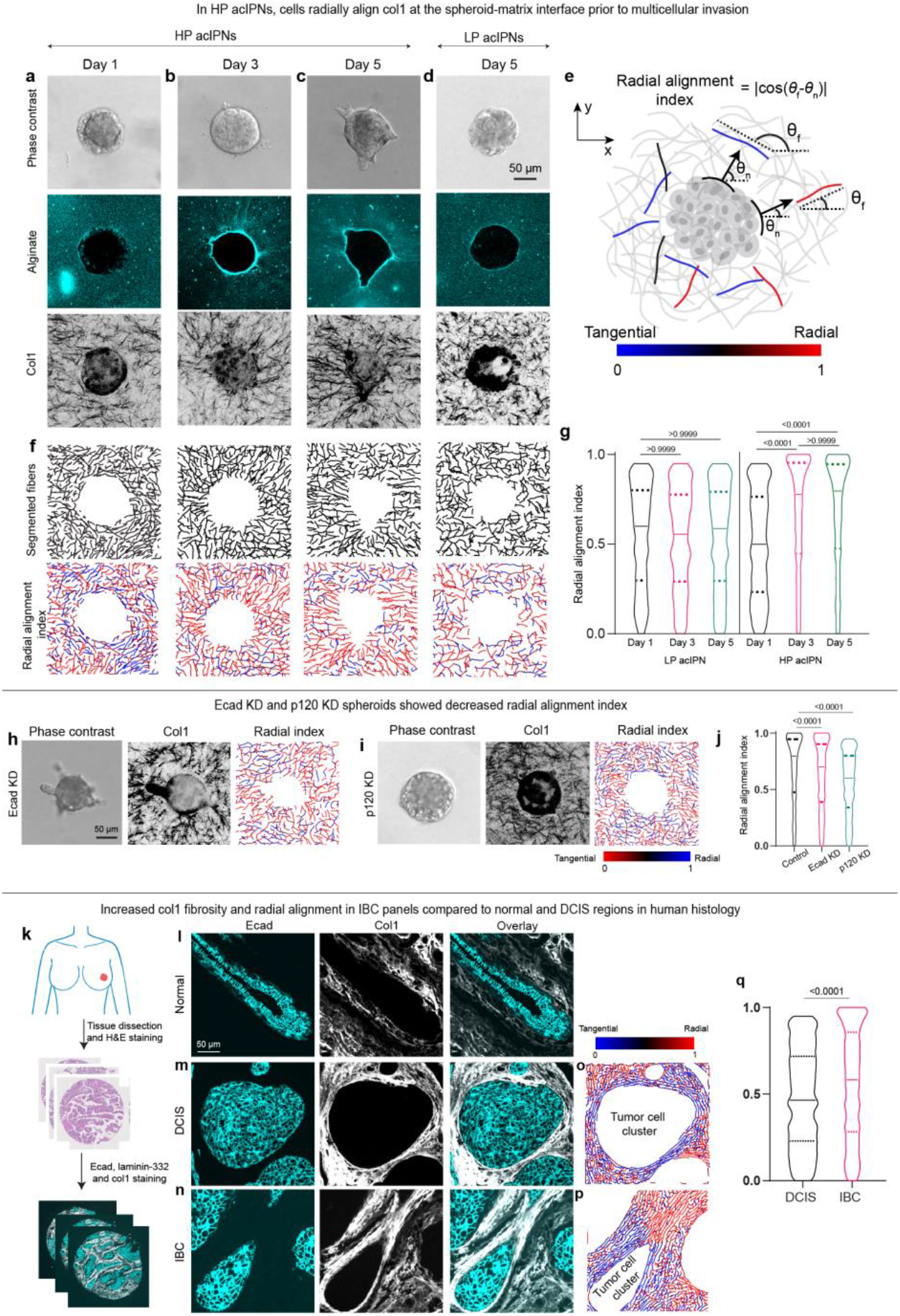
In stiff HP acIPNs, cells radially align col1 fibers prior to collective invasion and the radial alignment is similar to alignment observed in IBC. **a-d**, Phase contrast images of MCF10AT spheroids, and scanning confocal images of fluorescent alginate and col1 in the equatorial (or invasive) planes of the spheroids on days 1, 3 and 5 in HP acIPNs (**a-c**) and on day 5 (**d**) in LP acIPN. **e**, Schematic showing the computation of radial alignment index for each fiber segment with respect to the nearest normal vector of the spheroid surface. The dot product is computed based on the equation provided in the figure panel. Radially aligned fibers attain values of one (blue fibers) while tangentially aligned fibers are zeros (red fibers). **f**, Segmented fibers and the corresponding heatmaps of the radial alignment index of col1 in different conditions. **g**, Violin plot of radial alignment index for days 1, 3, and 5 in LP and HP acIPNs (n ≥ 6 spheroids; N = 3). **h-i**, Phase contrast images, confocal reflectance of col1 and the corresponding radial alignment index in Ecad KD and p120 KD conditions on day 5 in HP acIPN. **j**, Violin plots of radial alignment index in control, Ecad KD and p120 KD conditions (n ≥ 6 spheroids; N = 3). **k**, Schematic depicting the process of tumor dissection followed by H&E staining and then fluorescent staining for laminin, Ecad and col1. **l-n**, Fluorescent images of Ecad and col1 in normal (**l**), DCIS (**m**) and IBC (**n**) regions. **o-p**, Heatmap of radial alignment index of the col1 in panels **m** and **n. q**, Violin plot of radial alignment index of col1 in DCIS and IBC regions (n≥3 patients). Panels **g, j** – Kruskal-Wallis test with Dunn’s multiple comparisons. Panel **q** –Mann-Whitney test.

### Collective invasion involves proteases and requires cell-cell adhesions

We next sought to understand if cells use MMPs to invade, focusing on collective invasion in HP acIPNs at the BC-relevant stiffness of 3 kPa. Upon inhibition of MMPs using broad-spectrum protease inhibitors GM6001 and Marimastat, the average strand length of collectively invading cells in the HP acIPNs remained the same, but the strand width decreased (Extended Fig. 3). This is consistent with the previous findings that MMPs localize at the lateral sides of a collectively invading strand^14,20^. Nevertheless, collective invasion persisted. Thus, these studies indicate that MMPs support but are not strictly required for collective invasion.

**Figure 3:**
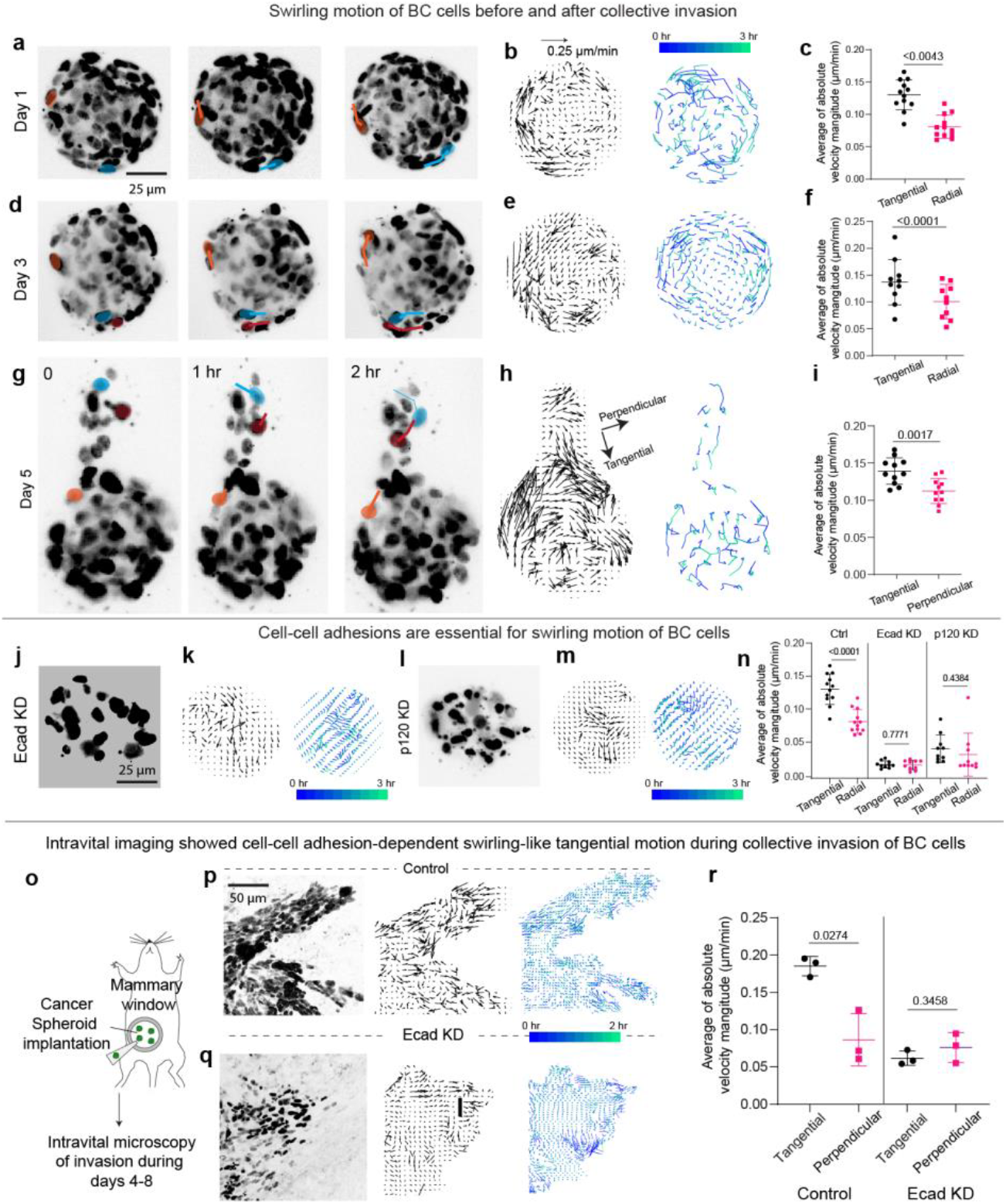
Cells exhibit tangential, or swirling, motion relative to the tumor-matrix interface. **a**, Confocal images of H2B-GFP nuclei in the equatorial plane of a MCF10AT spheroid at 0, 1 and 2 hrs on day 1. The highlighted cells were tracked over time. **b**, Instantaneous cell velocity field, and cell trajectories built over 3 hrs in the equatorial plane of MCF10AT spheroids on day 1. **c**, Average of absolute radial and tangential velocities near spheroid-ECM interface in the spheroid’s equatorial plane on day 1 (n ≥ 10 spheroids; N = 3). **d**, Confocal images of H2B-GFP nuclei in the equatorial plane of a MCF10AT spheroid at 0, 1 and 2 hrs on day 3. The highlighted cells were tracked over time. **e**, Instantaneous cell velocity field, and cell trajectories over 3 hrs of time lapse imaging in the equatorial plane of MCF10AT spheroids on day 3. **f**, Average of absolute radial and tangential velocities near spheroid-ECM interface in the spheroid’s equatorial plane on day 3 (n ≥ 10 spheroids; N = 3). **g**, Confocal images of H2B-GFP nuclei in the invasive plane of a MCF10AT spheroid at 0, 1 and 2 hrs on day 5. The highlighted cells are tracked over time. The highlighted cells were tracked over time. **h**, Instantaneous cell velocity field, and cell trajectories over 3 hrs of time lapse imaging in the invasive plane of MCF10AT spheroid on day 5. **i**, Average of absolute perpendicular and tangential velocities near spheroid-ECM interface in the spheroid’s invasive plane on day 5 (n ≥ 10 spheroids; N = 2). **j**, Confocal image of H2B-GFP nuclei in the equatorial plane of an Ecad KD MCF10AT spheroid on day 3. **k**, Instantaneous cell velocity field, and cell trajectories over 3 hrs of time lapse imaging in the equatorial plane of an Ecad KD spheroid on day 3. **l**, Confocal image of H2B-GFP nuclei in the equatorial plane of a p120 KD spheroid on day 3. **m**, Instantaneous cell velocity field, and cell trajectories over 3 hrs of time lapse imaging on day 3 of a p120 KD spheroid. **n**, Average of absolute radial and tangential velocities near spheroid-ECM interface in control, Ecad KD and p120 KD spheroids on day 3 (n ≥ 9 spheroids; N = 2). **o**, Schematic showing an orthotopic implantation of a BC spheroid in the mammary fat pad of a mouse and its intravital imaging between days 4 and 8 after the implantation. **p**, Confocal images of H2B-mCherry nuclei of collectively invading cells, instantaneous cell velocity field and the cell trajectories over 2 hrs of time lapse imaging of control (top) and Ecad KD (bottom) 4T1 cells. **q**, Average of absolute perpendicular and tangential velocities near tumor-stroma interface in control and Ecad KD conditions (n=3; N = 2 different mice). Panels **c, f, i, n, q** - Two-sided t-test.

Given that cells are collectively invading in HP acIPNs, we investigated the role of cell-cell adhesions, which regulate their collective behavior, focusing on Ecad and p120-catenin (p120), a non-redundant intracellular regulator of all cadherin-based adhesions. In HP acIPNs of 3 kPa, MCF10AT spheroids with shRNA-mediated knockdown (KD) of Ecad and p120 showed significant decrease in invasion compared to the control spheroids (Fig. 1r-s; Extended Fig. 4a-g). However, in HP acIPNs of 0.6 kPa, Ecad KD and p120 KD cells invaded at similar levels compared to the control cells, and the invasive strands mainly comprised a lane of single cells (Extended Fig. 4h-j). To complement these KD studies, MDA-MB-231 (MDA-MB) cells, an Ecad deficient (Ecad-) triple-negative BC cell, were induced to overexpress Ecad (Ecad+) (Extended Fig. 4k-n)^21^. Ecad+ MDA-MB spheroids invaded collectively compared to negligible invasion in Ecad-spheroids (Fig. 1s; Extended Fig. 4p-q). These data point to the essential role of cell-cell adhesions in collective invasion, though autonomous effects of Ecad and p120 could also play a role. Altogether, these data suggest that the collective action of cells, mediated by Ecad and p120, can overcome the barrier posed by BC-relevant stiff ECMs for invasion, provided the ECM exhibits sufficient mechanical plasticity (Fig. 1t).

**Figure 4:**
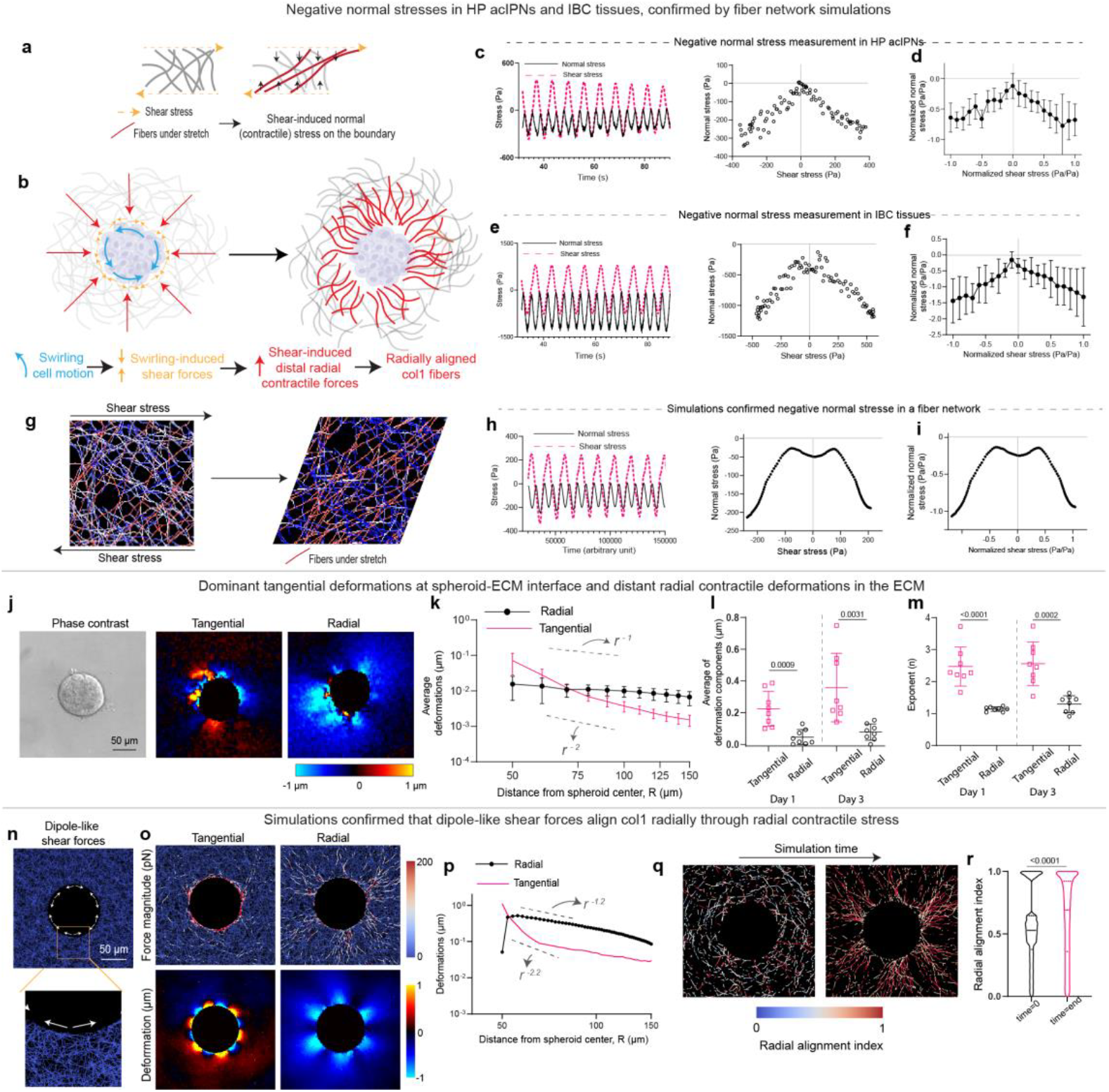
Negative normal stress in col1-rich ECMs mediates radial alignment of col1, with local shear stresses generated by cells leading to distal radial contractile stresses. **a**, Schematic depicting negative normal stress in a group of fibers, with shear stress leading to contractile or negative normal stresses generation by the stretched fibers. **b**, Schematic of the hypothesis: Shear stresses from tangential cell motion generate radially contractile stresses. The contractile stress radially aligns col1 fibers. **c**, Temporal plot of shear stress and normal stress during an oscillatory shear rheology of a HP acIPN at 20% strain amplitude, and the corresponding scatter plot. **d**, Average of normalized normal stresses with respect to normalized shear stress (n = 3). **e**, Temporal plot of shear stress and negative normal stress during shear rheology of an IBC tissue at 20% strain amplitude, and the corresponding scatter plot between the shear and the normal stress. **f**, Average of normalized normal stresses with respect to normalized shear stress (n = 4 patients). **g**, Snapshots of a fiber model before and after shear loading. Upon shearing, around 50% of the fibers undergo stretching (red fibers) and the other half are under compression (blue fibers). **h**, Temporal plot of shear stress and normal stress during oscillatory shear rheology of a fiber model at 20% strain amplitude, and the corresponding scatter plot between the shear and the normal stress. **i**, Scatter plot of normalized normal stresses with respect to normalized shear stress. **j**, Phase contrast image of MCF10T spheroid, and heatmaps of radial and tangential deformations in HP acIPNs on day 1. **k**, Line plots of the average of radial and tangential deformations with respect to the distance from center of MCF10AT spheroids on day 1 (n=8; N=3). **l**, Average of radial and tangential deformations near MCF10AT spheroid-ECM interface in HP acIPNs on days 1 and 3 (n=8; N=3). **m**, Exponents from power law fit of the line plots of radial and tangential deformations for days 1 and 3 (n=8; N=3). **n**, Snapshot of the fiber model with dipole-like shear forces applied at the inner circular boundary. **o**, Snapshots of force magnitude and deformations in tangential and radial directions. **p**, Line plot of the radial and tangential deformations with respect to the distance from the center of the circular geometry. The exponents from the power law fit of the tangential and radial line plots were 2.2 and 1.2 respectively. **q**, Snapshots of the radial alignment index of fibers that are being radially aligned at initial (left) and final (right) times of the simulation. Snapshots of the radial alignment index of all the fibers are provided in Extended Fig. 9a. **r**, Violin plot of the radial alignment index at initial and final times of the simulation. Panels **l, m -** Two-sided t-test. Panels **r –** Mann-Whitney test.

### Collectively invading cells align col1 radially prior to invasion

As cells collectively invaded into stiff HP acIPNs, the cells must make space for invasion by remodeling alginate and col1. To investigate this, we imaged fluorescently labelled alginate and col1 in LP and HP acIPNs on days 1, 3, and 5. While there was no apparent remodeling of alginate and col1 in HP acIPNs on day 1, on day 3, alginate was densified and col1 became radially aligned at the spheroid-ECM interface all around the spheroid even before the collective invasion (Fig. 2a-b). On day 5, a multicellular strand of collective cells was associated with a channel opening in the nanoporous alginate and with one of the radially aligned regions of col1 fibers at the spheroid-ECM interface (Fig. 2c). However, in LP acIPNs, alginate and col1 did not exhibit any remodeling (Fig. 2d, Extended Fig. 5a-b). The observation of radial alignment of col1 by day 3 in HP acIPNs was supported by quantification of radial alignment index^22^, which showed an increase in radial alignment index between day 1 and 3, but not much of a change in alignment between day 3 and day 5 (Fig. 2e-g). Upon cell lysis in LP and HP acIPNs, both alginate and col1 recoiled radially away from where the cells were, indicating a contractile-like stress state in the acIPNs (Extended Fig. 5c-f). Moreover, in the HP acIPNs, the alginate and col1 still remained partially remodeled, demonstrating that cells had permanently remodeled the ECM (Extended Fig. 5e).

**Figure 5:**
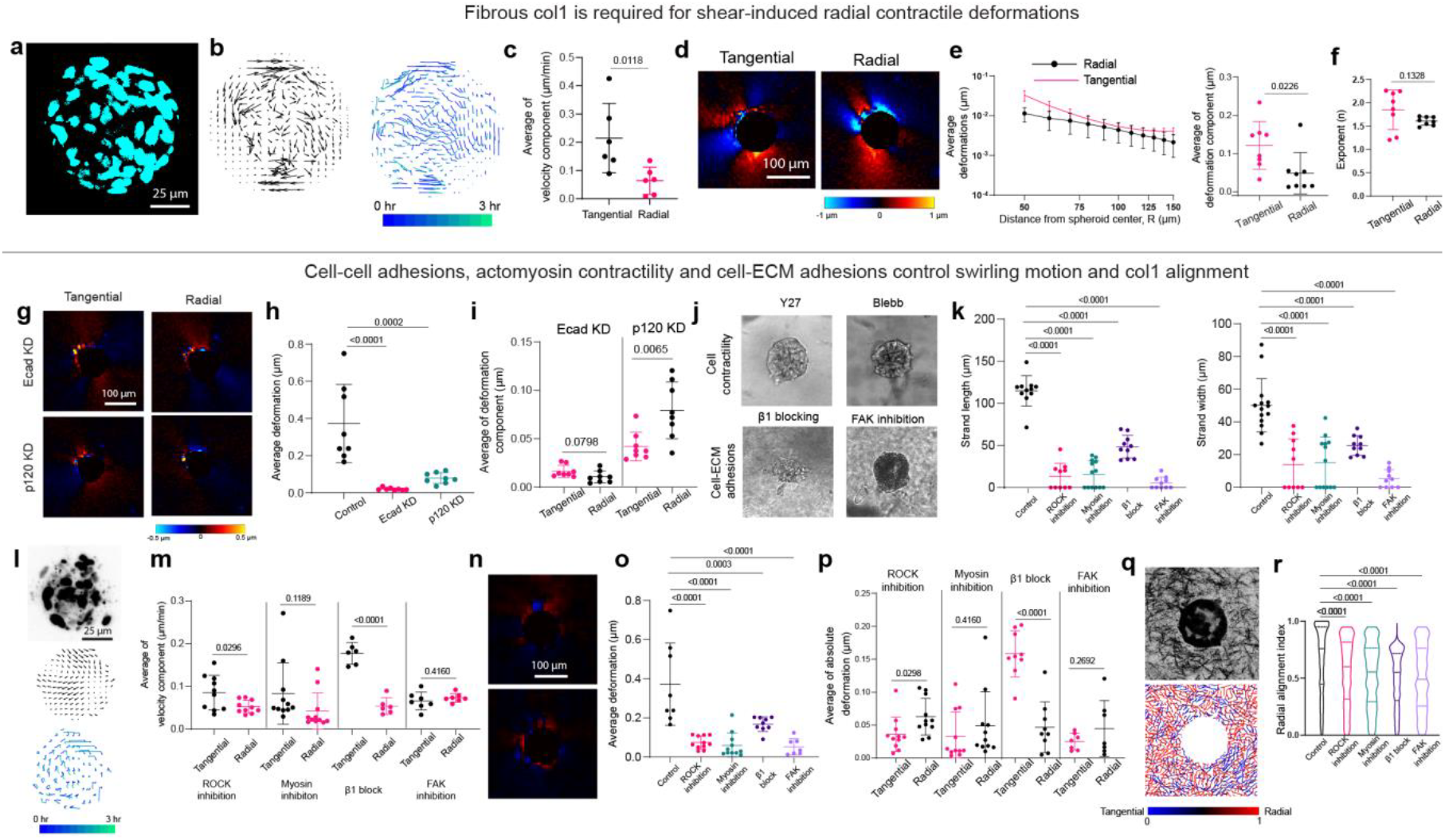
Col1 is necessary for negative normal stress-mediated radial stresses, and cell-cell adhesions, actomyosin contractility and cell-ECM adhesions mediate swirling motion and radial col1 alignment. **a**, Confocal image of H2B-GFP nuclei in the equatorial plane of MCF10AT spheroid. **b**, Instantaneous cell velocity field and cell trajectories over 3 hrs of time lapse imaging in the equatorial plane of a MCF10AT spheroid. **c**, Average of absolute radial and tangential velocity components near spheroid-ECM interface (n>5 spheroids in 2 hydrogels). **d**, Heatmap of tangential and radial deformation field on day 1. **e**, (Left) Line plots of the average of radial and tangential deformations with respect to the distance from center of MCF10AT spheroids. (Right) Average of radial and tangential deformations near MCF10AT spheroid-ECM interface (n=8; N=2). **f**, Exponents from power law fit of the line plots of radial and tangential deformations in panel **e** (n=8; N=2). **g**, Heatmaps of tangential and radial deformation fields around Ecad KD and p120 KD spheroids on day 3. **h**, Average deformation near spheroid-matrix interface in control, Ecad KD and p120 KD on day 3 (n>7; N=3). **i**, Average of radial and tangential deformations around spheroid-ECM interface in Ecad KD and p120 KD experiments (n=7; N=3). **j**, Phase contrast images of MCF10AT spheroids on day 5 in different conditions. **k**, Strand length and strand widths of collectively invading cells from MCF10AT spheroids on day 5 in different conditions (n>7; N=3). **l**, Confocal image of H2B-GFP nuclei (top), instantaneous cell velocity field (middle), and cell trajectories over 3 hrs of time lapse imaging (bottom) in the equatorial plane of ROCK-inhibited MCF10AT spheroid on day 3. **m**, Average of absolute radial and tangential velocity components near MCF10AT spheroid-ECM interface in different conditions on day 3 (n>5; N=3). **n**, Heatmap of tangential and radial deformation field around ROCK-inhibited MCF10AT spheroid on day 3. **o**, Average deformation near MCF10AT spheroid-ECM interface in different conditions (n>7; N=2). **p**, Average of radial and tangential deformations around spheroid-ECM interface in different conditions (n>7; N=2). **q**, Confocal reflectance of col1 and the corresponding heatmap of radial alignment index in ROCK-inhibited condition on day 3. **r**, Violin plot of radial alignment index in all conditions (n>3; N=2). Panels **c, e, f, i, m, p** - Two-sided t-test. Panels **h, k, o** - One-way ANOVA with Dunnett’s multiple comparisons. Panels **r** – Kruskal-Wallis test with Dunn’s multiple comparisons.

We then evaluated if the alignment of col1 is sufficient to facilitate collective invasion. Invasion assays were performed in LP acIPNs with locally aligned col1 domains, since cells did not invade in non-aligned LP acIPNs (Extended Fig. 5g-h). Spheroids invaded more in the LP acIPNs with pre-aligned col1 compared to the non-aligned condition, though not as much as in the HP non-aligned condition (Extended Fig. 5i-k). These demonstrate that alignment enables some degree of collective invasion in even matrices with low plasticity, but that increased matrix mechanical plasticity substantially enhances invasion.

Next, considering Ecad KD and p120 KD invaded significantly less in 3 kPa HP acIPNs, the roles of Ecad and p120 in mediating the radial alignment of col1 were examined. Both Ecad KD and p120 KD spheroids did not induce significant radial alignment of col1(Fig. 2h-j). Overall, these data show that collectively invading cells radially align col1 prior to the invasion in HP acIPNs in a cell-cell adhesion-dependent manner and that the aligned col1 then facilitates the collective invasion of cancer cells.

### Increased radial alignment of col1 at the tumor-stroma interface in IBC

Next, we investigated the physiological relevance of radial alignment of col1 by measuring col1 alignment at the tumor-stroma interface in normal, ductal carcinoma in situ (DCIS), and IBC regions of human BC tissues (Fig. 2k). Compared to normal and DCIS regions, IBC regions exhibited a loss of intact basement membrane (BM) component laminin-332 around Ecad+ cell clusters and a qualitative increase in fibrous col1 (Extended Fig. 6a-c). In DCIS regions, col1 around Ecad+ clusters was tangentially aligned with the tumor-stroma interface. However, IBC regions exhibited higher radial alignment at some parts of the boundary of the tumor cell cluster (Fig. 2m-r, q; Extended Fig. 6d), consistent with previously observed TACS-3 configuration in IBC^5,7–10^. Overall, these results establish an increase in radially aligned collagen, or TACS-3, at the tumor-stoma interface during the DCIS to IBC transition.

**Figure 6:**
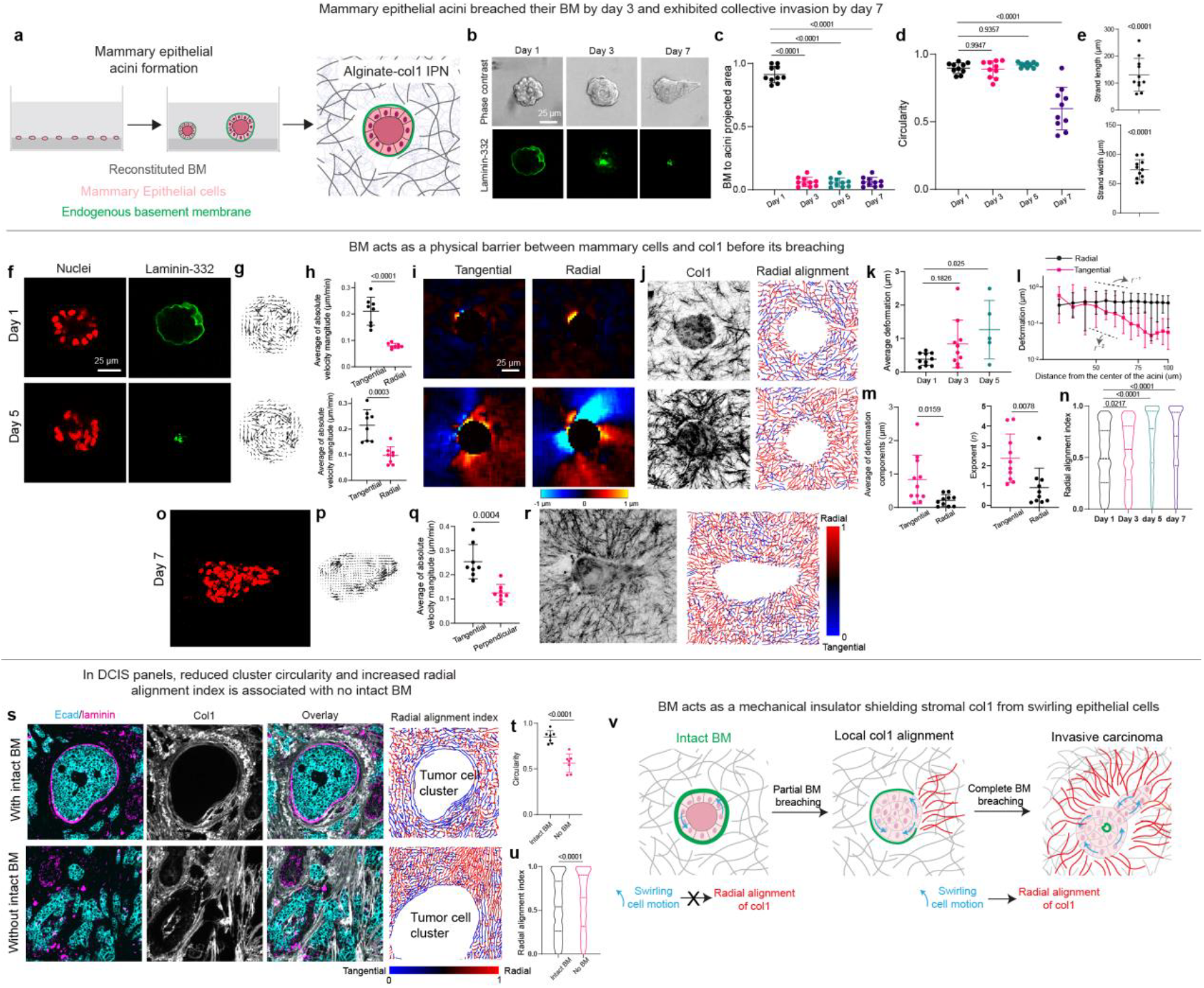
The basement membrane acts as a mechanical insulator, shielding the underlying stromal col1 matrix from swirling epithelial cells. **a**, Schematic of acini formation and its encapsulation in HP acIPNs. **b**, Representative phase contrast images and fluorescent images of laminin-332 on days 1, 3 and 7. **c-d**, Circularity and the ratio of BM to acini area of MCF10A acini on different days. **e**, Average strand width and length of collective invading strands from MCF10A acini on day 7 (n>9; N=3). **f**, Confocal images of H2B-RFP nuclei and laminin-332 in the equatorial plane of MCF10A acini on days 1 and 5. **g**, Vector field of instantaneous cell velocities on days 1 and 5. **h**, Average of absolute radial and tangential velocity components near the acini-ECM interface on days 1 and 5 (n>5; N=2). **i**, Heatmap of tangential and radial deformation field on days 1and 5. **j**, Confocal reflectance of col1 and the heatmap of radial alignment index of the confocal image on days 1 and 5. **k**, Average deformations at acini-matrix interface on days 1, 3, and 5 (n>9; N=2). **l**, Line plot of radial and tangential deformations with respect to the distance from the center of acini on day 5 (n>9; N=2). **m**, Average of radial and tangential deformations near acini-ECM interface on day 5, and the exponent of the power law fit of line plots of radial and tangential deformations in panel **l. n**, Boxplot of radial alignment index on days 1, 3, 5 and 7 (n>9; N=2). **o**, Confocal images of H2B-RFP nuclei in the invasive plane of MCF10A acini on day 7. **p**, Vector field of instantaneous cell velocities on days 1 and 5. **q**, Average of absolute radial and tangential velocity components near the acini-ECM interface on day 7 (n>5; N=2). **r**, Confocal reflectance of col1 and the heatmap of radial alignment index of the confocal image on day 7. **s**, Ecad, laminin-332, and col1 staining, and the heatmap of the radial alignment index of the col1 around tumor cell clusters with and without an intact BM. **t**, Circularity of Ecad+ tumor cell clusters with and without intact laminin (n=6 clusters; N=3 patients). **u**, Violin plot of radial alignment index of col1 around Ecad+ tumor cell clusters with and without an intact laminin (n>3 patients). **v**, A schematic showing how a no-breach, partial breach and a complete breach in the BM affect col1 remodeling. With a no-breach BM, the col1 is shielded from tangential cell motion. With a partial breach in the BM, cells remodel col1 locally through shear-induced radial contractile forces. With a complete breach of the BM, cells align col1 around the acini through shear-induced radial contractile stress. Panels **c, d, k -** One-way ANOVA with Dunnett’s multiple correction. Panel **e** - One-sample t-test. Panels **h, m, t** - Two-sided t-test. Panel **n -** Kruskal-Wallis test with Dunn’s multiple comparisons. Panel **u -** Mann-Whitney test.

### BC cells exhibit swirling motion before and during collective invasion

We next examined cell motion to understand the origins of TACS-3 realignment. Given that col1 was aligned radially by day 3 prior to the collective invasion in HP acIPNs, the cells at the spheroid-ECM interface are expected to exhibit radial motion to align col1 radially. In contrast to our expectation, timelapse imaging of nuclei on days 1 and 3, followed by quantitative analysis of both instantaneous cell velocities and cell trajectories, showed cells migrating tangentially to the interface in a non-coherent manner, which we describe as swirling, and exhibited no obvious outward or inward motion (Fig. 3a-f; Supplementary Video 2 - 3). Even after the collective invasion by day 5, both cells in the collectively invading strand and at the non-invasive front migrated tangentially to the interface in a swirling manner, despite the invading strand invading into the matrix (Fig. 3g-i; Supplementary Video 2-3). Consistent with this finding of a rotating group of cells leading invasion, leader cell marker cytokeratin 14 (K14) was expressed in cells in non-invasive regions in addition to the invasive front (Extended Fig. 7a)^23^. In LP acIPNs, swirling motion was also observed though there was no invasion (Extended Fig. 7b-j; Supplementary Video 4-5).

Given that Ecad and p120 were essential for the collective invasion and radial alignment of col1, we next investigated the motion of cells in Ecad KD and p120 KD spheroids. These cells did not exhibit any apparent tangential motion at the interface and were mainly stuck to their positions, pointing to the essential role of Ecad and p120-mediated cell-cell adhesions in swirling motion (Fig. 3j-n; Supplementary Video 6).

Next, we investigated the in vivo relevance of swirling motion by analyzing intravital imaging of collective invasion of 4T1 BC cells (control and Ecad KD) in an orthotopic mouse mammary model^6^ (Fig. 3o). Cell velocities and trajectories of a collectively invading strand of control 4T1 cells showed dominant tangential migration at the interface, mimicking the swirling motion on day 5 in HP acIPNs (Fig. 3p-q; Supplementary Video 7). In addition, Ecad KD 4T1 cells showed no apparent swirling motion (Fig. 3p-q; Supplementary Video 7). Altogether, intravital imaging of collective invasion in a BC mouse model confirms the in vivo relevance of the swirling motion of BC cells during collective invasion.

### Swirling motion induces distal radial contractile stress through negative normal stress

The observation of cells swirling at the spheroid-matrix interface and the simultaneous radial alignment of col1 is unexpected, as tangential cell motion would be expected to result in local shear stresses but not the radial contractile stresses that would be needed to radially align col1. This raises the question: how can the tangential motion of cells at the tumor-matrix interface lead to the radial alignment of col1?

A potential explanation for how local shear stresses can lead to radial alignment of col1 lies in the phenomenon of negative normal stress, which has been reported in fibrous ECMs such as col1 and fibrin^24^. When fibrous networks are sheared between two plates, the fibers undergo stretching and pull on the plates (or the plate pulls on the fibers), generating a negative (contractile) normal stress whose magnitude is comparable to the shear stresses (Fig. 4a). This is in contrast to materials like a polyacrylamide gel, which expand upon shearing and exhibit positive normal stress^24^. Therefore, we hypothesize that local shear stresses generated by the tangential motion of cells at the tumor-matrix interface result in radially contractile stresses due to negative normal stress, and such radial stresses in turn align col1 radially to facilitate invasion (Fig. 4b).

We first tested whether HP acIPNs, IBC tissues, and computational models of fibrous networks displayed negative normal stresses. HP acIPNs exhibited significant negative normal stresses when sheared and these stresses were comparable to the magnitude of applied shear stresses (Fig. 4c-d). Contrastingly, LP acIPNs exhibited both negative and positive normal stress, which are much lower than the applied shear stress (Extended Fig. 8a-c). Similar to HP acIPNs, IBC tissues also exhibited significant negative normal stresses that are comparable to the applied shear stresses, consistent with the fibrous nature of IBC tissue sections (Fig. 4e-f). Interestingly, in contrast to HP acIPNs and IBC tissues, healthy tissues exhibited positive normal stresses (Extended Fig. 8d-f), potentially because of their lower collagen fibrillarity (Extended Fig. 6a). Further, a computational fiber model confirmed the significant negative normal stresses upon shearing (Fig. 4g-i). Together, these establish negative normal stress as a key feature of many fibrillar col1-rich matrices, including high plasticity acIPNs and IBC tissues, with applied shear stresses leading to negative normal stress with comparable magnitude.

Next, we directly tested the hypothesis of whether negative normal stress could mediate radial alignment of col1 by measuring mechanical deformations of the matrix in the HP acIPNs. Radial and tangential deformations in the HP acIPN induced by tangential cell motion were measured, focusing on days 1 and 3. In HP acIPNs, tangential or shear deformations were localized to the spheroid-matrix interface and decayed rapidly with radial distance from the spheroid (Fig. 4j-k). The tangential deformations also exhibited alternating positive and negative values at the interface, suggesting a dipole-like shear forces coming from tangentially migrating cells. This is consistent with the understanding of contractile tension acting along the interface of cell-matrix interface induces a dipole-like deformation. Supporting this, the basal side of the cells at the basal surface exhibited actin fibers (Extended Fig. 8g). In contrast to the tangential deformations, the radial deformations showed lower deformations immediately adjacent to the spheroid but extended much further away from the spheroid in a radially inward contractile manner (Fig. 4j-k). This is consistent with the negative normal stress phenomenon and agrees with the acIPN recoil upon cell lysis noted earlier (Extended Fig. 5d, f). A power law fit (*u∼r*^-*n*^) of the radial and tangential deformations resulted in a smaller *n* for the radial deformations, demonstrating that the radial deformations propagated radially more distant than the tangential deformations (Fig. 4m). In LP acIPNs, while tangential deformations were localized at the spheroid matrix, both the radial and tangential deformations decayed over much shorter distances (Extended Fig. 8h-k).

Next, we tested our hypothesis of shear-induced radial alignment of col1 fibers using the fiber computational model (Fig. 4g). For this, the geometry mimicking the equatorial plane of cancer spheroids in HP acIPNs was created (Fig. 4n). At the inner circular boundary, dipole-like shear forces were applied to mimic the dipole-like shear deformations from the stress fibers (Extended Fig. 8g). Because of negative normal stress, shear forces at the boundary led to radially contractile forces distally from the boundary (Fig. 4o; Supplementary Video 8). These contractile forces generated radially contractile deformations which propagated over longer distances than the tangential deformations, mimicking the experimentally measured deformation profile (Fig. 4p). The radial contractile deformations eventually aligned the fibers radially, as quantified by the radial alignment index, thus confirming our hypothesis (Fig. 4q-r; Extended Fig. 9a). Simulations with other potential boundary conditions such as stochastic shear forces or uniform shear forces, did not capture the experimentally observed tangential and radial deformations (Extended Fig. 9b-e).

To establish the causal role of col1 in mediating shear-induced radial contractile stresses, we conducted invasion studies with tumor spheroids seeded in HP alginate-RGD hydrogels with no col1. In these ECMs, cells at the spheroid-ECM interface exhibited dominant tangential motion (Fig. 5a-c), resulting in dominant tangential deformations at the interface (Fig. 5d-e). However, the radial deformations decayed as fast as the tangential deformations, indicating there were no distal radial contractile stresses (Fig. 5e-f). Altogether, our data demonstrates that shear forces from swirling cell motion generate radial contractile stresses due to negative normal stress in col1-rich matrices with sufficient mechanical plasticity, and these radial contractile stresses align col1 radially.

### Cell-cell adhesions, cell contractility and cell-ECM adhesions control swirling motion and col1 radial alignment

We next examined the origins of the cell-induced shear stresses that lead to distal contractile radial stresses. Ecad KD and p120 KD spheroids did not generate significant ECM mechanical deformations, with no dominant tangential deformations. This is consistent with decreased invasion and radial alignment of col1 in those conditions (Fig. 5g-i).

At the single-cell level, for cells to generate forces, contractility-mediated actomyosin stress fibers at the spheroid-matrix interface are expected to generate dipole-like tangential forces. Upon inhibition of actomyosin contractility, via inhibition of myosin or ROCK, collective invasion was significantly decreased, with a significant decrease in swirling motion and matrix mechanical deformations, leading to no radial alignment of col1 (Fig. 5j-r; Extended Fig. 10a-c; Supplementary Video 9).

Cells apply contractile forces to col1 through col1-specific β1-containing integrins. Upon β1 integrin blocking, MCF10AT spheroids exhibited decreased invasion and diminished radial alignment, though they continued to exhibit swirling motion at a decreased magnitude (Fig. 5j-r; Extended Fig. 10a-d; Supplemental Video 11). Upon inhibition of focal adhesion kinase (FAK) which is involved in many integrin-based adhesions, MCF10AT spheroids showed decreased invasion with no swirling motion, decrease in ECM deformations, and col1 radial alignment (Fig. 5j-r Extended Fig. 10a-b, e; Supplemental Video 10). Overall, these studies establish that cell-cell adhesions, cell contractility and cell-ECM adhesions involving β1 integrin mediate swirling motion and induction of the TACS-3 col1 alignment that enables collective invasion.

### The BM insulates stromal col1 from swirling forces

In pre-invasive breast carcinoma, or DCIS, the laminin-rich BM separates the carcinoma cells from the surrounding col1-rich matrix (Extended Fig. 6b). Thus, we examined the role of BM in an established pre-invasive BC model involving mammary epithelial acini encapsulated in the HP acIPNs (Fig. 6a)^25^. In this model, the BM secreted by acini is comprised of a layer of laminin 332 on the inside, and type IV collagen on the outside, and further are induced to invade through the BM when the surrounding ECM is sufficiently stiff^25,26^. Indeed, upon encapsulating the acini in HP acIPNs of BC-relevant stiffness of 3 kPa, the mammary cells breached their BM by day 3 and invaded collectively into the HP acIPNs by day 7 (Fig. 6b-e).

On day 1, with an intact BM, the cells at the acini-BM interface exhibited dominant tangential motion, consistent with previous studies showing that cells in acini exhibit global rotation^27,28^ (Fig. 6f-h; Supplementary Video 12). Surprisingly, with the presence of the intact BM, the tangential cell motion did not generate any significant deformations on the surrounding col1-rich ECM or align col1 radially, indicating BM insulates the col1-rich ECM from the swirling motion (Fig. 6f-j; Supplementary Video 12). By day 5, the BM was breached, and the swirling cell motion generated significant local tangential deformations while long-range contractile-like radial deformations in the ECM aligning col1 radially, in agreement with our hypothesis (Fig. 6f-n; Supplementary Video 11). By day 7, while cells exhibited tangential motion at the acini-ECM interface, a collective strand of cells invaded along the radially aligned fibers (Fig. 6r).

Finally, we evaluated the relevance of TACS-3 and the concept of the BM as an insulator in human BC tissue specimens. Compared to tumors with an intact laminin layer, tumors without intact laminin showed a more invasive morphology, with decreased circularity and increased radial alignment of col1, in agreement with the acini measurements (Fig. 6s-u). Thus, the BM acts as a mechanical insulator, shielding the col1 rich matrix from remodeling forces prior to invasion. However, once the BM is breached, the carcinoma cells align col1 through shear-induced radial contractile stress to facilitate their invasion (Fig. 6v).

## Discussion

Here, we discover a mechanism of collective invasion whereby collectively invading cells radially align col1 fibers at the tumor-ECM interface to facilitate collective invasion in stiff ECM, provided that the ECM exhibits sufficient matrix mechanical plasticity. As cells swirl around at the spheroid-ECM interface, they generate local shear forces, which in turn lead to distal radial contractile forces due to the negative normal stress of col1-rich ECMs. These radial stresses, in turn, align col1 radially, facilitating the collective invasion of BC cells.

The TACS-3 signature, involving col1 fibers aligned radially at the tumor-matrix interface, associated with IBC was first reported in 2006, and has been shown to facilitate invasion by providing tracks for cells to migrate on. Further, TACS-3 has also been observed in skin melanoma and pancreatic adenocarcinoma^29,30^. The current paradigm is that the contractile activity of stromal CAF is responsible for inducing TACS-3^6,9,12,13^. Our mechanism explains how carcinoma cells can autonomously align col1 radially and could potentially act synergistically with CAFs.

A surprising aspect of the mechanism we report is that radial col1 alignment is driven by the swirling motion of cells. This is in contrast with the intuitive expectation that to radially align col1, cells would radially pull on the matrix, as a person might pull on a rope in a tug-of-war competition. Such swirling motion during the invasion suggests a group of leader cells, rather than a single K14+ leader cell^23^, and agrees with the emerging view of the collective front exhibiting cell-cell rearrangement^6,31–33^. The swirling motion observed here, in both in vitro studies and in orthotopic mouse models, resembles the tissue-scale rotational motion that has been observed during drosophila embryogenesis^34,35^ and in epithelial spheroids^27,28,36^, where it is implicated in BM maintenance, though the motion observed here is less coherent. These suggest the possibility that the swirling motion that is necessary for BM maintenance could have the negative effect of driving col1 realignment to facilitate invasion once the BM is lost. This is consistent with the pathological signature of the presence of BM as a distinguishing feature between DCIS and IBC^11,37^.

In our studies, enhanced matrix mechanical plasticity emerged as the key requirement for invasion. Conventionally, ECM remodeling is thought to be controlled by proteases. However, recent work has pointed towards the role of matrix mechanical plasticity in mediating permanent matrix remodeling^15,16^, affecting migration, invasion and mammary branching^16,17,38–41^. Matrix mechanical plasticity is distinct from stiffness, which has been a current parameter of focus in in vivo and in vitro models^1–4^. Matrix mechanical plasticity is also different from matrix viscoelasticity, which refers to viscous energy dissipation and a time-dependent mechanical response of a material in response to a stress or strain, though some of the molecular events that give rise to viscoelasticity also give rise to matrix mechanical plasticity (e.g. weak bonds)^42^. Our mechanical characterization of human tissues demonstrates that IBC tissues not only stiffen but also exhibit increased mechanical plasticity compared to their counterpart healthy tissues.

A key parameter that controlled the collectivity of invasion was matrix stiffness. Increasing evidence has indicated that cell-cell adhesions are necessary for early invasion and even metastasis despite the carcinoma cells undergoing partial epithelial-mesenchymal transition that could weaken cell-cell junctions^6,20,43–46^. Indeed, a recent study indicated that increased confinement in denser col1 gels promoted collective invasion over single-cell invasion^6,19^. Consistent with this finding, we find that at IBC-relevant stiffness, cells require cell-cell adhesions to invade. This suggests an interpretation that the collective action of cells is required to overcome the increased resistance of the stiffer matrix observed in IBC.

Finally, our proposed mechanism is based on the concept of negative normal stress, a fundamental phenomenon of fibrous matrices that was first described in 2006^24^. However, the physiological significance of negative normal stress was unclear and largely unstudied until now. Our rheological data showed that the IBC tissues exhibited greater negative normal stresses compared to healthy tissues, suggesting a possible role in BC pathogenesis. Moreover, the concept of negative normal stress could play a role during single-cell migration in a 3D confining fibrous matrix where the shear forces from the single cell can align col1 locally. Overall, the concept of negative normal stress in fibrous ECMs may have implications in other physiological processes during tissue development and cancer progression.

## Acknowledgements

Human tissue samples were obtained through the Stanford Tissue bank. We are grateful for receiving the tissues for science and thank all the human patients for donating their tissues. We thank Dr. Jake Song (Postdoctoral Researcher, Chaudhuri Lab) for providing fluorescent alginate and for the feedback on the manuscript. We acknowledge National Institutes of Health National Cancer Institute Grants R37 CA214136 and R01 CA290021 to O.C.; National Institute of General Medicine Sciences R35 GM136226 to L.H; and NIH U54 CA261694-01 to P.F. L.H. is an Irma T. Hirschl Career Scientist. The computational work was supported by National Institute of General Medical Sciences R01 GM151628 to T.K. We thank Prof. Valerie Weaver (University of California San Francisco) for helpful discussions.

## Author Contributions

Conceived and designed the experiments: A.S. and O.C. A.S. performed majority of the in vitro experiments with assistance from N.H.K.A. and L.H. Generation of MCF10AT KD cells: V.S. and Z.G. Intravital imaging: O.I. and P.F. Immunohistochemistry and imaging of human tissues: S.V and R.W. Designed and implemented the fiber computational model and wrote the corresponding results and methods section: M.F.R. and T.K. Wrote the manuscript: A.S. and O.C with input from all the authors.

## Data and code availability

All source data and codes will be provided for the final publication.

## Competing interests

The authors declare no competing interests.

## Materials and Methods

### Mechanical characterization of human tissues

Human healthy and IBC tissues were acquired from Stanford Tissue Bank. The tissues were stored in serum-free RPMI media at 4^°^C prior to mechanical characterization. Mechanical measurements were performed within two days of acquiring the samples. The mechanical testing regimen was as follows: two cycles of local indentation testing to measure matrix mechanical plasticity, shear rheology for shear and loss modulus, and amplitude sweep for negative normal stress measurements.

Tissue mechanical plasticity was measured using an Instron 5848 material testing system with a 1N load cell (Futek) or a DHR2 stress-controlled rheometer (TA instruments). Prior to indentation, the tissue samples were submerged in RPMI media at room temperature for 30 minutes. For indentation, a 1.5 mm spherical indenter was used, and the samples were indented up to 1 mm (all the samples were at least 4 mm in thickness) in depth at an indentation rate of 8 um/s in two cycles (Fig. 1e). In the first cycle, tissues were indented up to 1 mm in depth and the sample was unloaded by lifting the indenter. The sample was in this unloaded condition for one hour so that the sample can recover from any viscoelastic deformations. After 1 hour, in the second cycle, the sample was indented again for 1mm in depth at 8 μm/s. The difference in the indentation depths at 25% of the peak force in the first cycle was measured and was divided by the total indentation depth to compute mechanical plasticity. At least 3 different positions that were at least 3 mm apart were indented on each sample and the tissue mechanical plasticity was averaged over the different positions. After the indentation experiments, the tissue was unloaded and was left to sit at room temperature for 1 hour.

After the indentation experiments, the tissue mechanical stiffness was measured using the DHR2 stress-controlled rheometer (TA instruments). The tissues were punched into 8- or 6-mm discs (the disc sizes were similar between healthy and IBC tissues of the same patient). For the rheology, an 8 mm geometry top plate was used. To avoid any slip between the top or bottom plates and the tissue discs, sandpaper was attached to the bottom and top plates using a double-sided tape. After this, the tissue discs were placed between the sandpaper-attached plates and the top plate was lowered onto the tissue disc to a compressional pre-stress ranging between 200-600 Pa. This was done to avoid having any gap between the top plate and the tissue in case the tissue surface was not flat. Then, the storage and loss moduli were measured at 1 rad/s and 1% strain amplitude for 10 minutes, during which the moduli stayed consistently similar. After this, the tissue was unloaded and let the tissue sit in the RPMI media for 1 hour. The storage and loss moduli were averaged over the 10 minutes and the Young’s Modulus (E) was computed assuming a Poisson’s ratio (v) of 0.5 using the following equation:

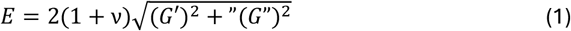

where *G*^*′*^ and *G*” are storage and loss moduli.

After measuring the storage and loss moduli, normal stresses applied by the tissues on the shear geometry were measured when the tissue discs were sheared using the 8mm geometry through a logarithmic strain amplitude sweep at 1 rad/s frequency. For this, the tissue discs were adhered to the top and bottom plates using superglue (Gorilla). Firstly, 8mm discs of kimwipe sheets were attached to the plates using a double-sided tape. Next, superglue was applied to the kimwipe sheet, and the tissue disc was placed right away between the plates. For better adhesion, the tissue disc was pre-stressed to 200-600 Pa, and the tissues were left to sit for 10 minutes for the superglue to act. Before starting the strain amplitude sweep, the normal force was set to zero. Then, the tissue was subjected to a strain amplitude sweep at 1 rad/s frequency with the amplitudes ranging from 0.1% to 100%. The positive and negative normal stress measurements were evaluated at 20% strain amplitude, as done previously^24^.

### Cell culture

Human MCF10A (ATCC) and MCF10AT cells (a gift from L. Wakefield, NIH) were cultured in Dulbecco’s modified Eagle’s medium/Nutrient Mixture F-12 (DMEM/F12) medium (Thermo Fisher) supplemented with 5% horse serum (Thermo Fisher), 20 ng/ml EGF (Peprotech), 0.5 ug/ml hydrocortisone (Sigma), 100 ng/ml cholera toxin (Sigma), 10 ug/ml insulin (Sigma) and 1% penicillin/streptomycin (Thermo Fisher). Both MCF10A and MCF10AT cells were stably transfected to express GFP/RFP-H2B. MDA-MB-231 and E-cadherin overexpressed MDA-MB-231 cells were a gift from P. Anastasiadis lab (Mayo clinic, Florida). 4T1 cells were a gift from S. Ponik lab (University of Wisconsin-Madison). MDA-MB-231 and 4T1 cells were provided with high glucose DMEM (Hyclone) with 10% fetal bovine serum (FBS) and 1% Pen/Strep. All cells were routinely split every 4-7 days using 0.05% trypsin/ethylenediaminetetraacetic acid (EDTA) and cultured in standard incubators at 37^°^C with 5% CO^2^.

### Generation of Ecad and p120 KD cells

All shRNA plasmids were generated as described previously^47^. Validated shRNA constructs for CDH1 (Ecad; TRC Clone TRCN0000237841) and CTNND1 (p120-catenin; TRC Clone ID TRCN0000333514) were cloned into an EF1a-GFP pSicoR plasmid downstream of the U6 promoter. Lentiviral packaging was performed by transfecting HEK294T/17 cells seeded in lentiviral packaging media with the plasmids for the gene of interest, pCMV-VSV-G (Addgene plasmid #8454) and pCMB delta R8.2 (Addgene plasmid #12263) using lipofectamine 3000 (ThermoFisher Scientific, #L3000015). The supernatant was harvested 24 and 48 hrs after transfection, centrifuged at 500 g for 10 minutes at 4^°^ C, and filtered using a 0.45 mm filter to remove debris. Next, the virus supernatant was incubated with Lenti-X concentrator (TakaraBio #631231) overnight and centrifuged at 1500 g for 45 min. The pellet was resuspended in 1/10^th^ of initial volume in DMEM/F12 medium, and the aliquots were stored at -80^°^ C. The lentiviral particle concentration was calculated by measuring the proportion of GFP+ or mCh+ by fluorescence activated cell sorting (FACS) 4-5 days after the infection.

A knockdown of Ecad and p120 proteins in MCF10AT cells was achieved by transducing the cells with the corresponding lentiviral particles at a multiplicity of infection of 2 for 40-60% transduction efficiency in the MCF10AT media containing 2 mg/mL polybrene (Millipore Sigma #TR1003). After 24 hrs, the virus-containing media was replaced with fresh media, and the cells grew to 80-90% confluence. Successfully transduced cells were sorted using FACS based on GFP+ expression. While these knockdowns were confirmed using western blots previously^47^, the knockdowns were further by looking at β-catenin localization at cell-cell junctions and coherent motion in cell monolayer migration (Extended Fig. 4a-c; Supplementary Video 12).

### Spheroid formation and MCF10A acini preparation

To generate spheroids of different cell lines, cells were trypsinized from tissue culture flasks and resuspended in pretreated Aggrewell multiwell plates (Aggrewell 400) to generate spheroids of 250 cells. The cells were incubated overnight which led to the formation of spheroids of 100 µm diameter. For spheroid formation of MCF10AT p120 KD cells, 2% w/v Matrigel was added to cell media so that the cells can aggregate together.

To generate MCF10A acini, six-well plates were coated with 120 µL of Matrigel (Corning) and the plates were placed at 37^°^C for 30 minutes. Then, single cell population of MCF10A cells at 40,000 cells/mL density was prepared in the growth media with 2% Matrigel in it. In each six-well plate, 2 ml of the solution was added. The media was replenished with the growth media with 2% Matrigel every two days. The single cells form acini within 5 days by secreting their endogenous BM, as confirmed previosly^25^. To encapsulate the acini in hydrogels, a 50 mM EDTA solution was added to the wells and the all the acini were extracted out by scraping the well surface. The acini-containing EDTA solution was incubated on ice for 20 min to disintegrate the Matrigel and was spun down for 5 min at 500xg for 5 min. After centrifugation, the supernatant was aspirated, and the acini was resuspended in the growth media. The live imaging of BM was performed by tagging the BM using Alexa Flour 488-conjugated laminin-5 antibody, clone D4B5 (Millipore Sigma). This was added to the resuspended medium at 1:400 ratio and incubated on ice for 1 hour. After the incubation, the media with the antibody was centrifuged again for 5 min at 500xg and the supernatant was removed to remove any excess antibody.

### Alginate-RGD preparation

The alginate-RGD was prepared in a two-step process, where step 1 involved lyophilizing the stock alginate and step 2 involved conjugating RGD peptide to the alginate chain. *Step 1:* High-molecular-weight (HMW; FMC Biopolymer, Protanal LF 20/40, 280 kDa) and low molecular weight ultrapure (LMW; Pronovo VLVG) sodium alginate was dialyzed against deionized water using dialysis tubing with a molecular weight cutoff of 3.5 kDa. The dialyzed alginate was subsequently treated with charcoal, sterile filtered and lyophilized. *Step 2*: To prepare HMW or LMW-RGD alginate, RGD peptides were coupled to the alginate using carbodiimide chemistry. The lyophilized alginate (both HMW and LMW) from Step 1 was dissolved in 0.1 M 2-(N-morpholino)ethanesulfonic acid (MES), 0.3 M sodium chloride buffer at pH 6.5 at 1% w/v for 24 hours. After 24 hours, 274mg of N-hydroxysulfosuccinimide (Thermo Fisher Scientific), 484 mg of N-(3-3-dimenthylaminopropyl)-N’-ethylcarbodiimide hydrochloride (Sigma Aldrich), and 112 mg of glycine-glycine-glycine-gycline-arginine-glycine-aspartate-serine-proline (GGGGRGDSP) (Peptide 2.0) peptide were sequentially mixed, and the reaction was allowed to happen for 20 hours. This yields a 1500 µM RGD concentration in a 2%w/v alginate solution, or 20 RGD peptides on one alginate chain on average. Next, both the alginates were dialyzed against deionized water for 2-3 days with a 3.5 kDa dialysis tubing, filtered with activated charcoal, sterile filtered, lyophilized and then reconstituted in serum-free DMEM at 3.5 % w/v solution.

### Mechanical characterization of alginate-collagen IPNs

All alginate-collagen IPNs (acIPNs) were formulated to attain a 4.8 mg/mL alginate and 1.6 mg/mL collagen, but with different molecular weights and calcium concentrations to attain different stiffnesses and their corresponding plasticity conditions (Supplementary Table 1). First, rat tail collagen-1 (Corning) was mixed with 10X DMEM in 1:10 ratio and the pH was adjusted to 7.4 using 1 M sodium hydroxide (NaOH), and was placed on ice for 15 minutes. Next, in a 1.5 mL Eppendorf tube, the stock alginate (3.5% w/v) solution, serum-free DMEM and the neutralized col1 were mixed (volumes as defined in Supplementary Table 1) and pipetted into a 1 mL luerlock syringe (Cole-Parmer). In a different syringe, DMEM and CaSO4 were added (volumes as defined in Table 1). Both the syringes were now mixed using a female-female luerlock coupler (Value Plastics) at least 4 times and then the gel was deposited onto a substrate of interest. The calcium in the alginate has been previously shown to not affect cell behavior^48^.

The shear rheology of acIPNs was performed using a 25 mm geometry at 1 rad/s and 1% shear strain using the AR2000EX stress-controlled rheometer (TA instruments). Before making the gel, PLL-coated coverslips were attached to the top and bottom plates of the rheometer using a double-sided tape to adhere the hydrogel to the plates and avoid any slip. Once acIPNs of required formulation was mixed using the syringe system, the hydrogel was deposited onto the PLL-coated coverslip attached to the bottom plate. Immediately, while the hydrogel was curing, the top plate was lowered to a gap of 500 um. A gel volume of 500 μL approximately forms a complete disk with 25 mm in diameter and 500 µm in height. Next, mineral oil (Sigma) was applied to the edges of the geometry or the hydrogel to prevent dry-out. Then, the shear and loss moduli were measured at 1 rad/s and 1 % shear strain for 8 hours. The moduli usually equilibrated in the last 2 hours. The average of these two moduli was taken in the last two hours and the Young’s modulus was computed, as described in the human tissue characterization section. Using the same setup with 25 mm geometry with PLL-coated coverslips, the normal stresses were measured when the hydrogel was sheared using the 25 mm geometry through a logarithmic strain amplitude sweep at 1 rad/s frequency. The positive and negative normal stress measurements at 20% strain amplitude were considered, as done for the tissue sections.

Mechanical plasticity was measured similar to the measurements of human tissues, as described in the human tissue characterization section. Once acIPNs of required formulation were mixed using the syringe system, the hydrogel was deposited into a 35-mm petri dish (Celvis) and let the gel sit for 1 hour to crosslink completely. Once the gel was crosslinked, the hydrogel was hydrated using DMEM. Next, the two-cycle indentation was performed using the 1.5 mm spherical indentation using the Instron 5848 material testing system with a 1N load cell or the AR2000EX stress-controlled rheometer. Mechanical plasticity was computed, as described in the tissue characterization section. At least 3 different positions that were at least 5 mm apart were indented on each sample, and mechanical plasticity were averaged over the different positions.

### Quantification of single and collective invasion in spheroid assays

The invasion of cancer spheroids was quantified using the following five metrics: circularity, number of collectively invading strands, average strand length, average strand widths, and single cell invasions. Acini circularity indicates breakage of spherical symmetry as the invasion occurs, with lower circularity indicating greater invasion. All these metrics were quantified either from the phase contrast, or fluorescent F-actin and nuclei (DAPI) imaging of the spheroids. The circularity index was measured in ImageJ by manually drawing an outline in the equatorial plane of the spheroid boundary. The number of collectively invading strands, their lengths and widths, and single cell invasions were measured manually from the phase contrast or fluorescent images of F-actin in an invading spheroid.

### Confocal microscopy

Confocal microscopy was performed with a laser-scanning Leica SP8 confocal microscope or a Nikon Ti2-E inverted microscope, both fitted with a temperature and incubator control suitable for live imaging (37 °C, 5% CO_2_). A 10x 0.45-NA dry objective or 20x 0.75-NA dry objective was used for the Nikon microscope. A 10× 0.4-NA dry or 25x 0.95-NA objective was used for the Leica microscope. For imaging col1, confocal reflectance setting was used in the Leica SP8 confocal microscope using a 488 nm laser.

### Pharmacological inhibitions

The concentrations of pharmacological inhibitors were as follows: 25.0 μM blebbistatin (Abcam, myosin II inhibitor), 10.0 μM FAK inhibitor (Selleck Chem), 25.0 μM GM6001 (Millipore, MMP inhibitor), 100.0 μM marimastat (Tocris, MMP inhibitor), and 25.0 μM Y-27632 (Y27) (Sigma, ROCK inhibitor). β1 integrin blocking was performed using 1μg/mL β1 anti-integrin antibody clone P5D2 (Abcam, #ab24693). Appropriate controls used include IgG from mouse serum (Sigma, #I5381). Media containing pharmacological inhibitors or functional antibodies was added on day 1 after spheroid encapsulation and was added whenever the media was replaced (days 3 and 5). Invasion was quantified towards the end of day 5.

### Immunostaining cells in bulk hydrogel

Hydrogels were rinsed with PBS and fixed with 4% paraformaldehyde for 1 hr. The fixed hydrogels were rinsed with PBS 10 mM EDTA to enable diffusion of antibodies. Next, the hydrogels were incubated at 4^°^ C overnight in PBS containing 1mM calcium, 0.5% Triton and 3% bovine serum (blocking buffer) to permeabilize cells and block non-specific binding respectively. After this, the hydrogels were incubated with primary antibodies at 4^°^ C overnight in the blocking buffer. After the primary antibody incubation, the hydrogels were rinsed and were incubated with secondary antibodies overnight. Primary antibodies used were anti-E-cadherin (BD Biosciences, #610181; 1:100), Vimentin (Santa Cruz Biotechnology, #sc-6260; 1:200), and cytokeratin 14 (ThermoFisher, #MA%-11599; 1:200). Secondary antibodies used were Alexa Fluor 555 goat anti-mouse IgG_1_ (#<u>A21240</u>, 1:1,000) and Alexa Fluor 555 goat anti-rabbit (#<u>A21244</u>, 1:1,000). For nucleus, either the transfected H2B-GFP or HOECHST 3342 (ThermoFisher; #62249; 1:1000) was used. F-actin was stained with SiR-actin (Cytoskeleton Inc., #CY-SC001; 0.1 nM).

### Col1 fiber characterization and radial alignment index

To measure the radial alignment index of individual col1 fibers in our in vitro confocal scanning imaging, firstly, col1 fibers were segmented using CTFire^22^ from the confocal reflectance images of the col1. This segmentation provides the vertices of each segmented fiber, using which the angle of the orientation of each fiber was computed by fitting a straight line to the fiber (*θ*_*f*_). Secondly, the boundary of the cancer spheroids or the mammary acini was outlined manually using ImageJ, and the angle (*θ*_*n*_) of the normal vector at each pixel on the boundary was computed. Lastly, for each of the segmented fiber, the nearest normal vector of the tumor boundary was found using KNN function in Matlab and computed the absolute of the dot product using the following equation, which quantifies the radial alignment index for the fiber:

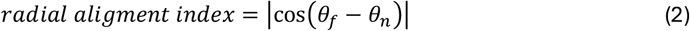

### Measuring cell velocities and trajectories

To quantify cell velocities, fast iterative digital image correlation (FIDIC)^49^ was employed between two consecutive confocal images, 15 or 30 minutes apart, of H2B-GFP imaging of the equatorial planes and the invasive planes in cancer spheroids and the acini. The resulting displacements were divided by the time interval to compute velocities. For the resolution, subset sizes of 64×64 or 32×32 pixels using a spacing of 16 or 8 pixels, which resulted in a spatial resolution of approximately 3 μm.

To compute radial and tangential cell velocities in spheroids that were circular in cross sections, the cell velocities vectors were transformed into a polar coordinate system with the center of the spheroid as the origin. In the case of the invasive spheroid or the acini, firstly, the boundary of the spheroid (or the acini) was outlined and the tangential and normal vectors of the boundary were computed at each pixel of the boundary. Next, for each pixel on the boundary, the nearest cell velocity vector was found using KNN function in Matlab (2022). Lastly, the dot product between the tangential (normal) vector of the boundary and the nearest cell velocity vector was computed to determine the tangential (perpendicular) velocity components. To compute the average of absolute radial and tangential velocities at the spheroid-matrix interface, the absolute of radial (and tangential) velocities was taken within one cell length (∼20 µm) from the spheroid boundary and then averaged over the boundary for each spheroid or acini.

Cell trajectories were built using cell displacements computed over a time lapse of 3 hours, as described previously^50^. Briefly, the calculation was performed starting with an initial evenly spaced grid with the distance between the grid points matching an approximate cell length. With these initial grid points, displacements between two consecutive time points were interpolated to each grid point, and the interpolated values are added to the coordinates of each grid point to give an estimate of each cell’s position at the next time point. This was repeated over the time lapse of 3 hours, thereby creating a list of points for each cell corresponding to that cell’s trajectory.

### Measuring ECM mechanical deformations

All the experiments involving measuring mechanical deformations in the ECM were performed by embedding 0.5 µm fluorescent beads (carboxylate modified; Life Technologies) at 4% (v/v) concentration in the acIPNs. To quantify the deformations, time-lapse imaging of the fluorescent beads in the ECM was performed at 15-minute time intervals around the equatorial plane of the cancer spheroids or the mammary acini prior to the collective invasion. Next, FIDIC was employed between the consecutive time lapse images using 32 × 32-pixel subsets centered on a grid with a spacing of 8 pixels (∼1.6 μm). These mechanical deformations were deformations induced by the cells over time.

To plot radial and tangential deformations, the cartesian coordinate system was transformed into a polar coordinate system with the spheroid or the acini center as the origin. To plot the average of radial (or tangential) deformations with respect to the distance from the center of the spheroid, the average of the radial (or tangential) deformations at a radial increment of every 5 um was computed, and all the deformations within +/-2.5 um at each of the radii were considered for the average. At each of the radii, temporal averaging of radial and tangential deformations was performed. For plotting the deformations on a log-log scale, since the radial deformations are mostly contractile (or negative), the deformations were multiplied by a negative sign. For tangential deformations, the average of the absolute tangential deformations with respect to each radius was measured. To avoid any artifacts in the FIDIC measurements within the spheroid, the spheroid boundary was manually outlined and deformations outside this boundary were only considered.

### Tissue immunohistochemistry

*Immunostaining and imaging:* Paraffin-embedded tissue microarray (TMA) sections (Stanford TA 417, 419 and 424 – normal, DCIS and IBC regions respectively) were stained for Immunofluorescence (IF). Primary antibodies used for IF staining include laminin-332 (Santa Cruz Biotech catalogue no. sc-28330) conjugated with ArgoFluor676, E-cadherin (Rarecyte, conjugated with ArgoFluor546) and collagen-I (Abcam ab138492 Anti-Collagen I antibody [EPR7785]) conjugated with ArgoFluor706. Followed IF staining protocol recommended by Rarecyte. The slides were imaged using Rarecyte (Orion Imager).

*Measuring circularity index in DCIS tissue microarray sections*: In each panel of the DCIS TMA (Stanford TA 419), Ecad-positive clusters with and without laminin were identified based on laminin-332 fluorescent imaging, and an outline was drawn around the clusters to measure their circularity in ImageJ.

*Measuring radial alignment index in DCIS and IBC TMA*: In each panel of the DCIS and IBC TMAs, regions containing clusters of Ecad+ cells and the col1 around the clusters alone were cropped. Similar to col1 analysis of our in vitro imaging, col1 fibers were segmented using ctFIRE and the boundary of the Ecad+ cell clusters was outlined to get the boundary. Based on this boundary, the radial alignment index of the segmented col1 fibers was computed based on equation (2) as described in the corresponding section. A box plot was plotted to show the distribution of the radial alignment index for each panel corresponding to a different patient (Extended Fig. 6d). For the main panel, the data was from all the patients for DCIS and IBC regions was combined and plotted as a violin plot (Fig. 2q).

### Intravital imaging of collective invasion of 4T1 cancer cells in a mouse orthotopic model

*Intravital imaging of collective invasion:* Protocol for 4T1 cancer spheroid implantation in mouse mammary fat pad and the intravital imaging of the collective invasion has been established and described in detail previously^6^.

Briefly, Balb/c female mice (6-8 weeks old; Charles River Laboratories) were anesthetized and 4T1 H2B-mCherry cell spheroids (control and Ecad KD) were implanted in the 4^th^ mammary fat pad. Intravital imaging was performed on a customized upright TrimScope II multiphoton microscope (LaVision BioTec, Miltenyi Biotec) equipped with three tunable Ti:Sa lasers (Coherent, Ultra I and II) and an Optical Parametric Oscillator (Coherent/APE) using a ×20 Olympus XLUMPlanFI ×20/0.95 NA water-immersion objective with a custom-made objective heater (37 °C) and up to 5 photomultipliers (Hamamatsu, Alkali H6780-01, H6780-20 or GaAsP H7422A-40) with 593/40 nm or 620/60 nm (mCherry).

### Theoretical model

To mimic the experimental system of col1 surrounding a spheroid, a 3D matrix between two cylindrical boundaries was created. The outer and inner radii of the cylindrical boundaries are 200 μm and 50 μm, respectively, and their height is 1 μm. The matrix consists of fibers interconnected by cross-linkers. At the beginning of simulations, the matrix is self-assembled via the nucleation and polymerization of fibers and the binding of cross-linkers to pairs of fibers. The cross-linkers can unbind from the fibers in a force-dependent manner. The endpoints of the fibers near two cylindrical boundaries bind to the boundaries irreversibly. After matrix assembly, while the endpoints bound to the outer boundary remain relatively stationary, those bound to the inner boundary undergo clockwise or counterclockwise rotation, depending on imposed conditions, to apply shear forces to the matrix. More details of the model are described below, and key parameter values are listed in Supplementary Table 2.

#### Simplification of fibers and cross-linkers and Brownian dynamics

Fibers consist of serially connected cylindrical elements. Cross-linkers comprise two cylindrical elements connected at their center point. The endpoints of all the cylindrical elements have x, y, and z positions. The velocities of all the endpoints at each time point are calculated via the Langevin equation with inertia neglected:

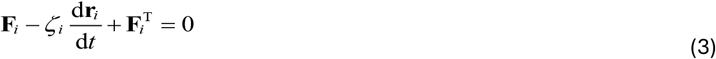

where **r**_i_ is a position vector of the ith endpoint, ζ_i_ is a drag coefficient, t is time, **F**_i_ is a deterministic force, and 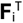 is a stochastic force satisfying the fluctuation-dissipation theorem^51^:

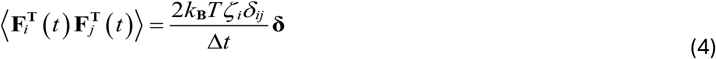

where δ_ij_ is the Kronecker delta, **δ** is a second-order tensor, and Δt = 4.0×10^−4^ s is a time step. The drag coefficients of cylindrical elements are calculated using an approximated form for a cylindrical object^52^.

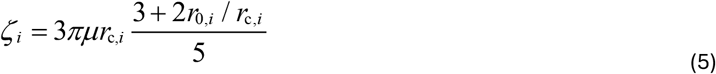

where μ is the viscosity of surrounding medium, and r_0,i_ and r_c,i_ are the length and diameter of the cylindrical element, respectively. The positions of all the endpoints are updated using the calculated velocities, d**r**_i_/dt, and the Euler integration scheme:

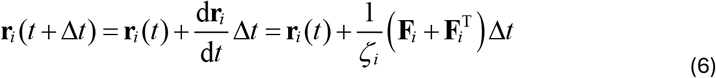

Deterministic forces include extensional forces maintaining equilibrium lengths and bending forces maintaining equilibrium angles. The extensional and bending forces for fibers and the extensional force for cross-linkers originate from harmonic potentials:

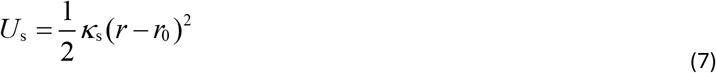

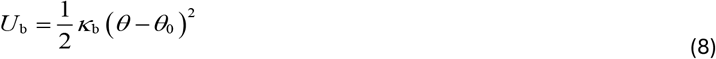

where *k*_*s*_ and *k*_*b*_ are extensional and bending stiffnesses, r and r_0_ is instantaneous and equilibrium lengths of cylindrical elements, and θ and θ_0_ are instantaneous and equilibrium angles formed by adjacent elements. An equilibrium length of fiber elements (r_0,f_ = 1.0 μm) and an equilibrium angle formed by two adjacent fiber elements (θ_0,f_ = 0 rad) are regulated by extensional (*κ*_*s,f*_) and bending stiffnesses of the fibers (*κ*_*b,f*_), respectively. An equilibrium length of cross-linker arms (r_0,xl_ = 20 nm) is maintained by extensional stiffness (*κ*_*s,xl*_). Forces exerted on the fiber elements by bound cross-likers are distributed onto two endpoints located of the fiber elements.

#### Dynamics of fibers and cross-linkers

The formation of fibers is initiated by the appearance of one cylindrical element whose length is 1.0 µm. The element is elongated by the addition of identical cylindrical elements. Average fiber length is 1 µm. There is no disassembly or depolymerization for the fibers.

Cross-linkers bind to binding sites located every 100 nm on fibers with the rate of *k*_+,xl_ and also unbind from fibers with a force-dependent rate, *k*_−,*xl*_, governed by Bell’s law^53^:

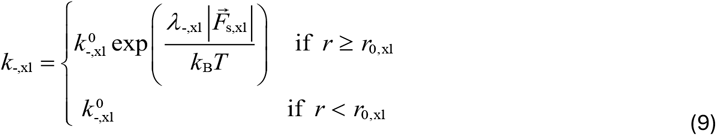

where 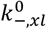 is the zero-force unbinding rate, λ_-,xl_ represents sensitivity to applied force, *k*_*B*_*T* is thermal energy, and 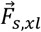 is a vector for a spring force acting on a cross-linker element. Only when the spring force is a tensile force, *k*_−,*xl*_ increases beyond its base rate, 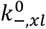.

#### Boundary conditions

The endpoints of fibers located within 1 μm from the outer cylindrical boundary are permanently bound to the boundary at the rate of k_+,bnd_ and constrained via a spring with the stiffness of κ_s,bnd_ until the end of simulations. The endpoints of the fibers located within 1 μm from the inner cylindrical boundary are also bound to the boundary at the rate of k_+,bnd_. The binding points of those fibers bound to the inner boundary rotate clockwise or counterclockwise. There are three different ways to determine the direction of the rotation. First, in the dipole way, the rotation direction is clockwise if a remainder after dividing the initial circumferential angular position of the binding point (measured counterclockwise from the +x axis) by 60^°^ is smaller than 30^°^ and counterclockwise if the remainder is equal to or greater than 30^°^. Second, in a stochastic way, the rotation direction is selected between clockwise or counterclockwise at equal probability at the moment of binding. Third, in the uniform way, the rotation direction is always counterclockwise. The binding point of the fibers is rotated in the specific direction by 0.001^°^ at each time point until the fibers feel a tensile force greater than 100 pN.

The two cylindrical boundaries and two z boundaries keep all the matrix elements within a 3D annular region by exerting repulsive forces to elements crossing the boundaries. The magnitude of the repulsive forces is directly proportional to an overlap distance with the boundaries and its strength defined by κ_r,bnd_. The direction of the repulsive forces is always normal to the boundaries.

## Statistical analysis

Unless otherwise noted, all measurements were done in at least 3 biologically different samples (n; ex: spheroids) and in at least two biological replicates (N; ex: hydrogels) from separate experiments. Each figure legend indicates n and N and the corresponding statistical tests performed. For human tissue data, the number of patients (n) is mentioned in the legend. Statistical analyses were performed using GraphPad Prism (version 10.1.0; https://www.graphpad.com/). *P* < 0.05 was considered to be statistically significant, and the P values are reported until 4^th^ decimal point. Error bars in all the plots, except for the violin plots, indicate mean +/-standard deviation. In the violin plots, the top and bottom lines represent the interquartile range, and the middle line represent the median.

## Supplementary Video captions

**Supplementary Video 1:** Time lapse phase contrast imaging of MCF10AT cancer spheroids in HP acIPNs of different stiffness over different days.

**Supplementary Video 2:** Time lapse confocal imaging of H2B-GFP nuclei in the equatorial (days 1 and 3) or invasive (day 5) planes of MCF10AT cancer spheroids in 3 kPa HP acIPNs.

**Supplementary Video 3:** Time lapse confocal imaging of H2B-GFP nuclei and col1 (reflectance) in the equatorial planes of MCF10AT cancer spheroids in 3 kPa HP acIPNs on days 1, 3 and 5.

**Supplementary Video 4:** Time lapse confocal imaging of H2B-GFP nuclei in the equatorial (days 1 and 3) or invasive (day 5) planes of MCF10AT cancer spheroids in 3 kPa LP acIPNs.

**Supplementary Video 5:** Time lapse confocal imaging of H2B-GFP nuclei and col1 (reflectance) in the equatorial planes of MCF10AT cancer spheroids in 3 kPa LP acIPNs on days 1, 3 and 5.

**Supplementary Video 6:** Time lapse confocal imaging of H2B-RFP nuclei in the equatorial planes of Ecad KD and p120 KD MCF10AT spheroids in 3 kPa HP acIPNs on day 3.

**Supplementary Video 7:** Intravital imaging of H2B-mCheery of control and Ecad KD 4T1 cells in vivo 4 days after multicellular spheroid implantation in the mouse mammary fat pad.

**Supplementary Video 8:** Evolution of force profile in the fiber model with dipole-like shear forces at the circular boundary.

**Supplementary Video 9:** Time lapse confocal imaging of H2B-RFP nuclei in the equatorial planes of Y27-treated and blebbistatin-treated MCF10AT spheroids in 3 kPa HP acIPNs on day 3.

**Supplementary Video 10:** Time lapse confocal imaging of H2B-RFP nuclei in the equatorial planes of MCF10AT spheroids treated with integrin β1 blocking functional antibody and FAK inhibitor in 3 kPa HP acIPNs on day 3.

**Supplementary Video 11:** Time lapse confocal imaging of H2B-mCherry, laminin-332 and col1 (reflectance) of MCF10A acini on days 1, 3 and 5 in 3 kPa HP acIPNs.

**Supplementary Video 12:** Time lapse imaging of the monolayer of H2B-GFP control cells, cytoplasmic mCherry Ecad KD and cytoplasmic mCherry p120 KD MCF10AT cells.

**Supplementary table 1:** Hydrogel composition for different stiffness and their corresponding plasticity conditions.

| Stiffness | Plasticity | Syringe 1 |  |  | Syringe 2 |  |
| --- | --- | --- | --- | --- | --- | --- |
| | | Alginate<br>($\mu\text{L}$)<br>(3.5 %w/v) | DMEM<br>($\mu\text{L}$) | Col1 ($\mu\text{L}$)<br>(3.2 mg/mL;<br>Neutralized) | DMEM | CaSO4 (488<br>mM) |
| 0.6 kPa | LP (HMW) | 107 | 180 | 300 | 72 | 3 $\mu\text{L}$ |
| | HP (LMW) | 107 | 180 | 300 | 70 | 5 $\mu\text{L}$ |
| 1.2 kPa | LP (HMW) | 107 | 180 | 300 | 66 | 9 $\mu\text{L}$ |
| | HP (LMW) | 107 | 180 | 300 | 63 | 12 $\mu\text{L}$ |
| 3 kPa | LP (HMW) | 107 | 180 | 300 | 59 | 16 $\mu\text{L}$ |
| | HP (LMW) | 107 | 180 | 300 | 55 | 20 $\mu\text{L}$ |

**Supplementary table 2:** List of parameters employed in the agent-based computational model.

| Symbol | Definition | Value |
| --- | --- | --- |
| $r_{0,f}$ | Length of fibers | $1.0 \times 10^{-6}$ [m] |
| $r_{c,f}$ | Diameter of fibers | $1.0 \times 10^{-7}$ [m] |
| $\theta_{0,f}$ | Bending angle formed by adjacent fibers | 0 [rad] |
| $\kappa_{s,f}$ | Extensional stiffness of fibers | $4.0 \times 10^{-3}$ [N/m] |
| $\kappa_{b,f}$ | Bending stiffness of fibers | $8.27 \times 10^{-20}$ [N·m] |
| $r_{0,xl}$ | Length of a cross-linker arm | $2.0 \times 10^{-8}$ [m] |
| $r_{c,xl}$ | Diameter of a cross-linker arm | $1.0 \times 10^{-8}$ [m] |
| $\kappa_{s,xl}$ | Extensional stiffness of cross-linkers | $2.0 \times 10^{-3}$ [N/m] |
| $k_{+,xl}$ | Binding rate of cross-linkers | $1.0 \times 10^2$ [$\mu\text{M}^{-1} \cdot \text{s}^{-1}$] |
| $k_{-,xl}^0$ | Zero-force unbinding rate of cross-linkers | $1.0 \times 10^{-6}$ [$\text{s}^{-1}$] |
| $\lambda_{-,xl}$ | Force sensitivity of cross-linker unbinding | $4.0 \times 10^{-10}$ [m] |
| $C_f$ | Fiber density | $\sim 2.85$ [fiber/$\mu\text{m}^3$] |
| $\langle L_f \rangle$ | Average length of fibers | $\sim 1$ [$\mu\text{m}$] |
| $R_{xl}$ | Ratio of # of cross-linkers to # of fibers | $\sim 5.72$ [cross-linker/fiber] |
| $k_{+,bnd}$ | Binding rate of fibers to cylindrical boundaries | $1.0 \times 10^3$ [$\mu\text{M}^{-1} \cdot \text{s}^{-1}$] |
| $\kappa_{r,bnd}$ | Strength of repulsive forces from boundaries | $1.69 \times 10^{-3}$ [N/m] |
| $\kappa_{s,bnd}$ | Extensional stiffness for boundary binding | $4.0 \times 10^{-3}$ [N/m] |
| $\Delta t$ | Time step | $4.0 \times 10^{-4}$ [s] |
| $\mu$ | Viscosity of medium | 8.6 [kg/m·s] |
| $k_B T$ | Thermal energy | $4.142 \times 10^{-21}$ [J] |

## Notes

### Competing Interest Statement

The authors have declared no competing interest.

