## Extended Figures for "Swirling motion of breast cancer cells radially aligns collagen fibers to enable collective invasion"

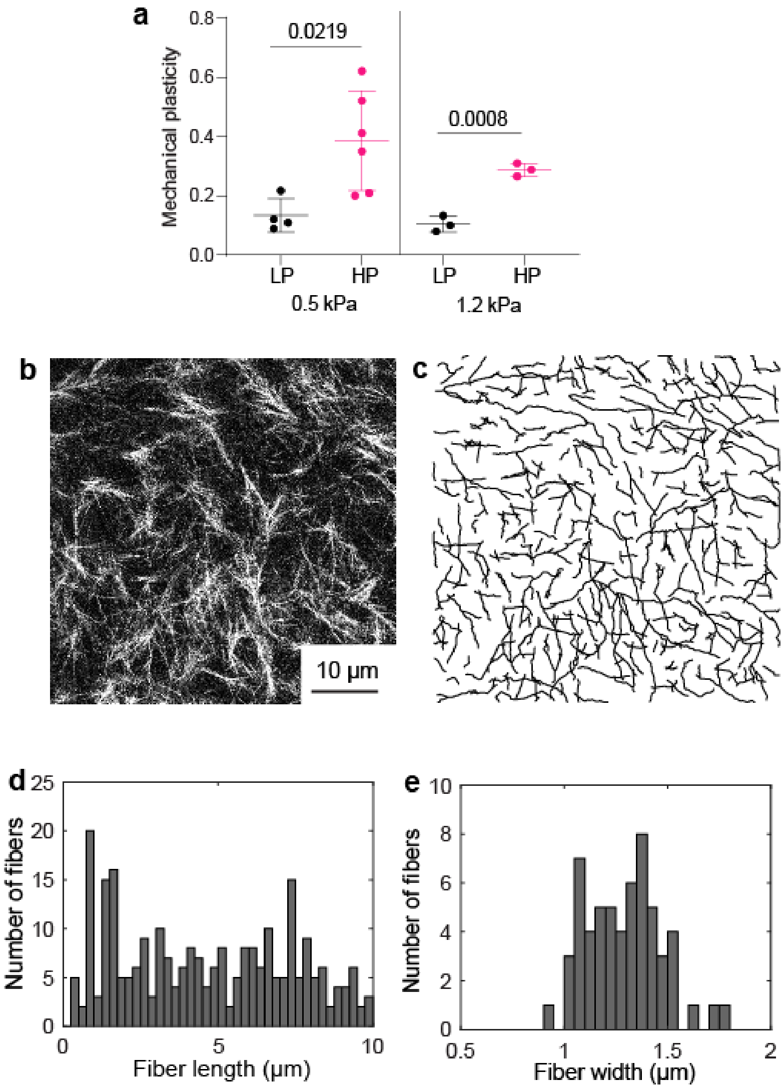


**Extended Figure 1: Mechanical plasticity and measuring col1 fiber length and width in acIPNs. a,** Mechanical plasticity of LP and HP acIPNs at 0.6 kPa and 1.2 kPa (n≥3 different hydrogels). **b-c,** Confocal reflectance of col1 and its segmented image. **d-e,** Histogram of fiber length and fiber width of col1 fibers in panel **b**. Panel **a** – Two-sided t-test.

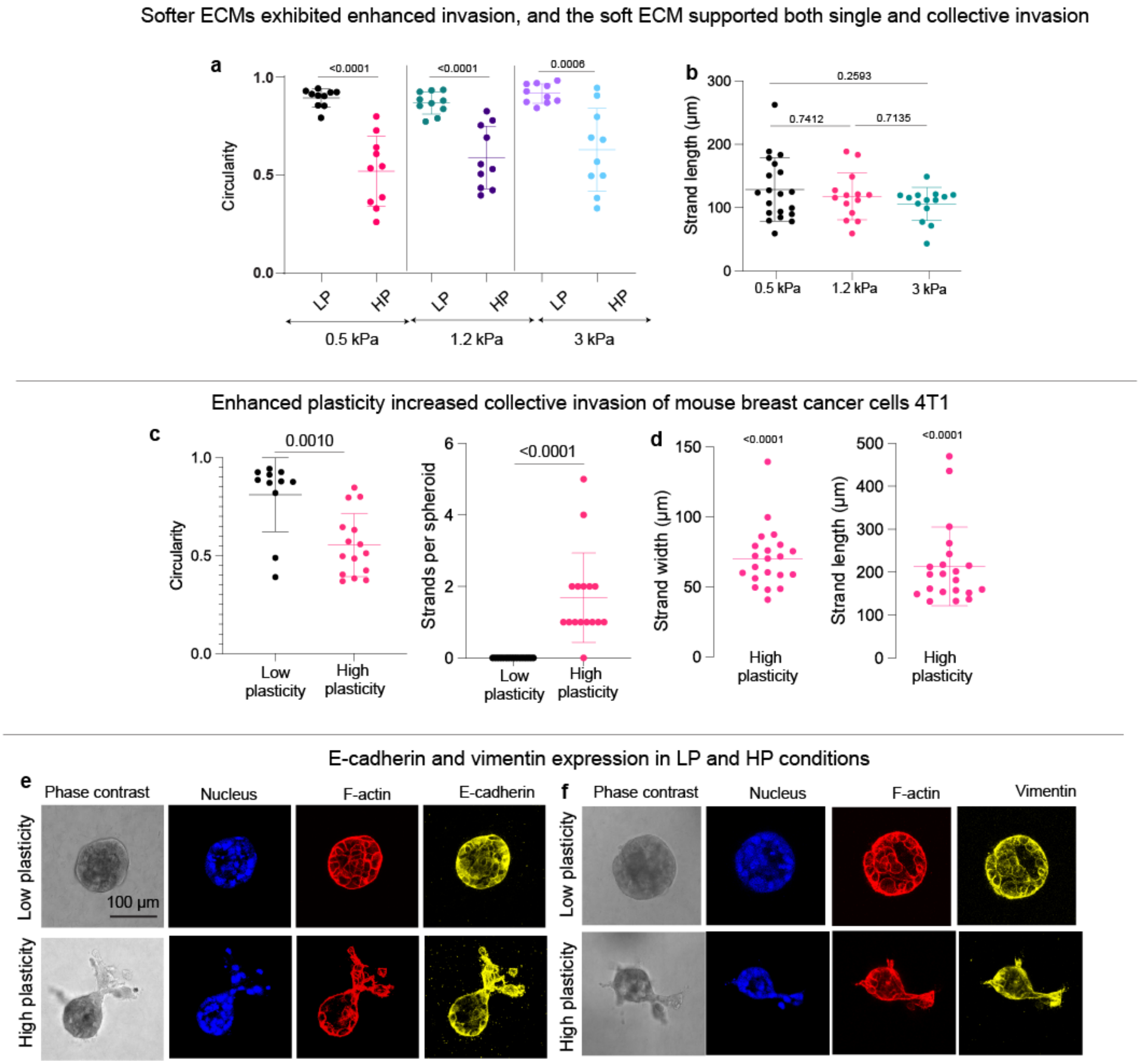


**Extended Figure 2: Collective invasion of MCF10AT spheroids in different stiffness conditions and of 4T1 cells, and Ecad and Vimentin expression in MCF10AT spheroids.** **a,** Circularity of spheroids in LP and HP acIPNs of 3 different stiffness conditions (n≥ 9 spheroids; N=3). **b,** Average strand width of collectively invading strand in HP acIPNs of 3 different stiffness conditions (n≥ 9 spheroids; N=3). **c-d,** Circularity, strands per spheroid and their average strand with and lengths of 4T1 spheroids in LP and HP acIPNs at 3 kPa stiffness (n≥ 9 spheroids; N=2). **e-f,** Phase contrast images, and scanning confocal images of DAPI, F-actin, E-cadherin and Vimentin in the equatorial (or invasive) plane of MCF10AT spheroids on day 5 in LP and HP acIPNs of 3 kPa stiffness. Panels **b** – One-way ANOVA with Tukey's correction for multiple comparisons. Panels **a, c** – Two-sided t-test. Panel **d -** Wilcoxon signed-rank test.

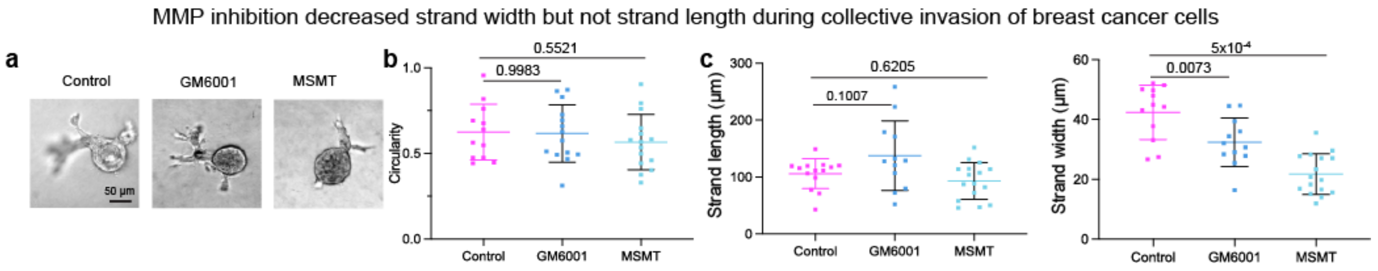


**Extended Figure 3: MMP inhibition on collective invasion of MCF10AT spheroids. a,** Phase contrast images of MCF10AT spheroids on day 5 in control, GM6001-treated and marimastat-treated conditions. **b-c,** Circularity, average strand length and strand widths of collectively invading cells of MCF10AT spheroids in control, GM6001-treated and marimastat-treated conditions (n≥ 9 spheroids; N=3). Panels **b, c** – One way ANOVA with Dunnett's multiple comparisons.

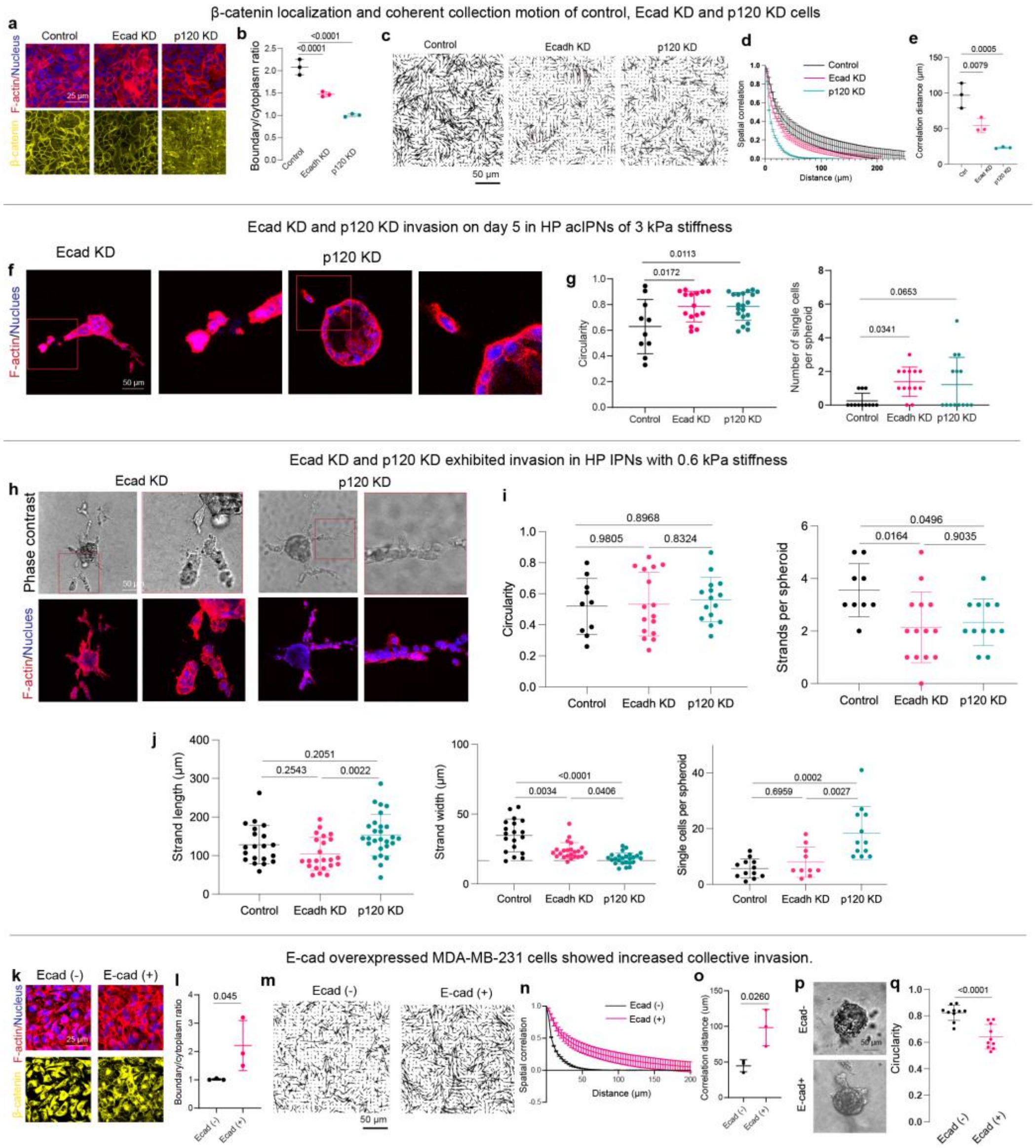


**Extended Figure 4: β-catenin expression and spatial velocity distribution in monolayers of control and cell-cell adhesion-perturbed cells, and their collective invasion in LP and HP acIPNs of 0.6 kPa and 3 kPa. a,** Confocal images of F-actin, nuclei and β-catenin in monolayers of control, Ecad KD and p120 KD cells. **b**, Ratio of average intensity of β-catenin at cell boundary to cytoplasm in monolayers of control, Ecad KD and p120 KD cells (n=3; N=1). **c,** Instantaneous velocity field of cell migration in monolayers of control, Ecad KD and p120 KD cells. **d,** Spatial correlation of the velocity field in monolayers of control, Ecad KD and p120 KD cells. **e,** Distances at which the spatial correlation dropped to 0.2 in monolayers of control, Ecad KD and p120 KD cells (n=3; N=1). **f,** Confocal scanning images of F-actin and nuclei and the zoomed in versions in the equatorial planes of Ecad KD and p120 KD spheroids on day 5 in HP acIPNs at 3 kPa stiffness. **g,** Circularity and number of single cells per spheroid in control, Ecad KD and p120 KD spheroids in HP ECMs at 3 kPa stiffness (n≥ 9; N=3). **g,** Phase contrast images, and scanning confocal images of F-actin and nuclei and the zoomed in versions in the invasive planes of Ecad KD and p120 KD spheroids on day 5 in HP ECMs at 0.6 kPa. **i,** Circularity and number of collectively invading strands per spheroid in control, Ecad KD and p120 KD spheroids in HP acIPNs of 0.6 kPa stiffness (n n≥ 9; N=3). **j,** Average strand length and width, and single cells per spheroid in control, Ecad KD and p120 KD spheroids on day 5 in HP acIPNs at 0.6 kPa stiffness (n≥ 9; N=3). **k,** Confocal images of F-actin, nuclei and β-catenin in monolayers of Ecad- and Ecad+ MDA-MB-231 cells. **l**, Ratio of average intensity of β-catenin at cell boundary to cytoplasm in Ecad- and Ecad+ MDA-MB-231 monolayers (n=3; N=1). **m,** Instantaneous velocity field of cell migration in monolayers of Ecad- and Ecad+ MDA-MB-231 cells. **n,** Spatial correlation of the velocity field in monolayers of Ecad- and Ecad+ MDA-MB-231 cells. **o,** Distances at which the spatial correlation dropped to 0.2 in monolayers of Ecad- and Ecad+ MDA-MB-231 cells. Each data point corresponds to the field of view of a monolayer (n=3; N=1). **p,** Phase contrast images of Ecad- and Ecad+ spheroids on day 5 in HP acIPNs at 3 kPa stiffness. **q,** Circularity of Ecad- and Ecad+ MDA-MB-231 spheroids on day 5 in HP acIPNs at 3 kPa stiffness (n=9; N=2). Panels **b, e, g, i, j** – One way ANOVA with Dunnett's multiple comparisons when compared to control and with Tukey's correction for multiple groups. Panels **l**, **o, q** – two-sided t-test.

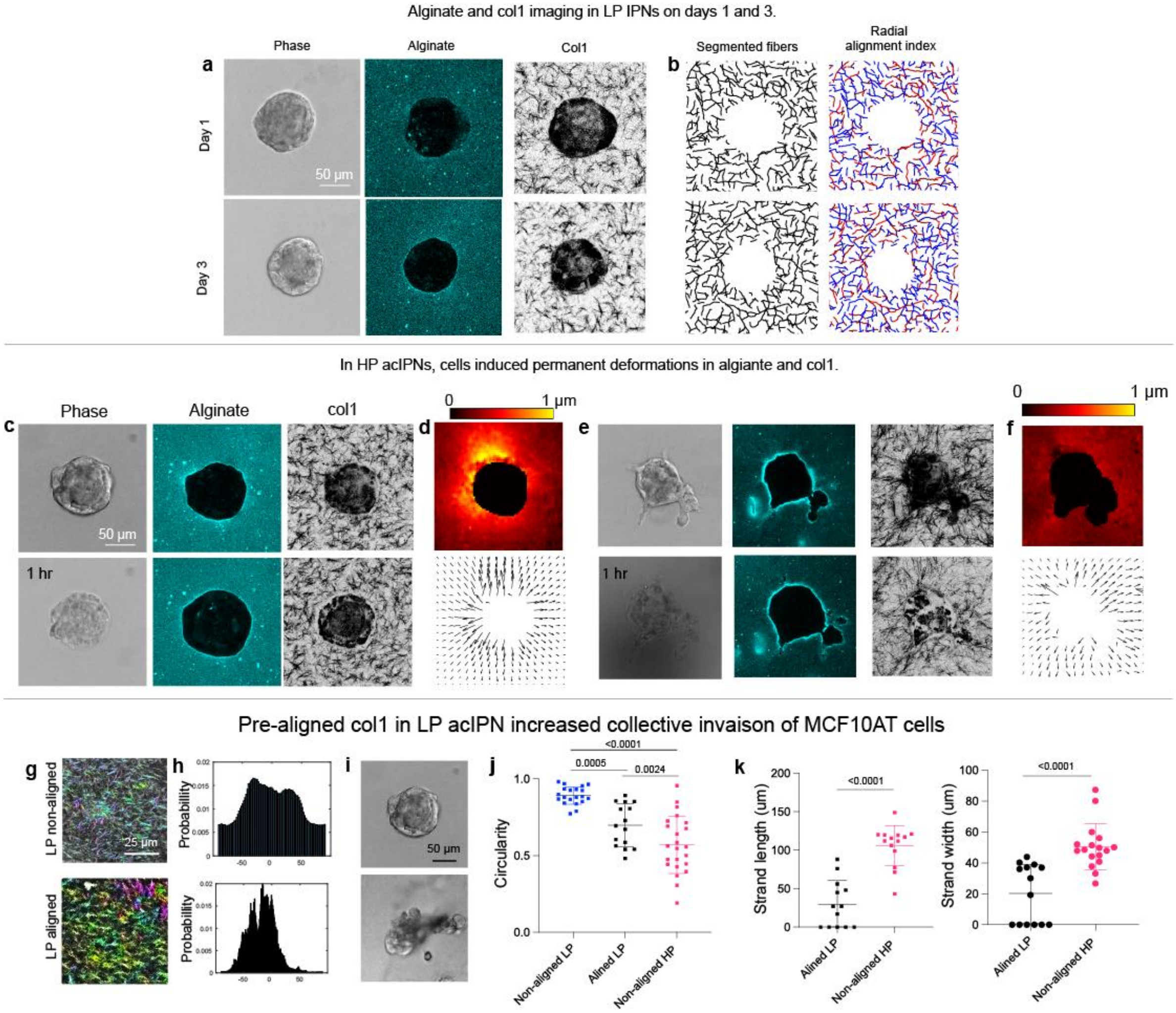


**Extended Figure 5: Radial alignment index in LP acIPNs, cell lysis, and invasion in LP acIPNs with pre-aligned col1. a,** Phase contrast images of MCF10AT spheroids, and confocal scanning images of alginate and col1 in the equatorial plane of MCF10AT spheroids on days 1 and 3 in LP acIPNs. **b,** Segmented col1 fibers and their radial alignment index of the col1 images in panel **a**. **c,** Phase contrast images of MCF10AT spheroids, and scanning confocal images of alignate and col1 in the equatorial planes of MCF10AT spheroids before (top) and 1 hr after (bottom) cell lysis in LP acIPN on day 5. **d,** Heatmap (top) and vector field (bottom) of ECM deformation 15 min after cell lysis in LP acIPN on day 5. **e,** Phase contrast images of MCF10AT spheroids, and scanning confocal images of alignate and col1 in the equatorial planes of MCF10AT spheroids before (top) and 1 hr after (bottom) cell lysis in HP acIPN on day 5. **f,** Heatmap (top) and vector field (bottom) of ECM deformation 15 min after cell lysis in HP acIPN on day 5. **g,** Confocal reflectance imaging of col1 in LP acIPNs with non-aligned and pre-aligned col1. **h,** Histogram of the angles of col1 fibers in non-aligned and pre-aligned conditions with respect to horizontal axis. **i,** Phase contrast images of MCF10AT spheroids in LP acIPNs with non-aligned and pre-aligned col1. **j,** Circularity of MCF10AT spheroids on day 5 in LP acIPNs with non-aligned and pre-aligned col1 and in HP acIPNs (n≥9 spheroids; N=2). **k,** Average strand length and strand width in LP acIPNs with non-aligned and pre-aligned col1 (n ≥ 9; N=2). Panel **j** – One-way ANOVA with Tukey's correction for multiple comparisons. Panel **k** - Two-sided t-test.

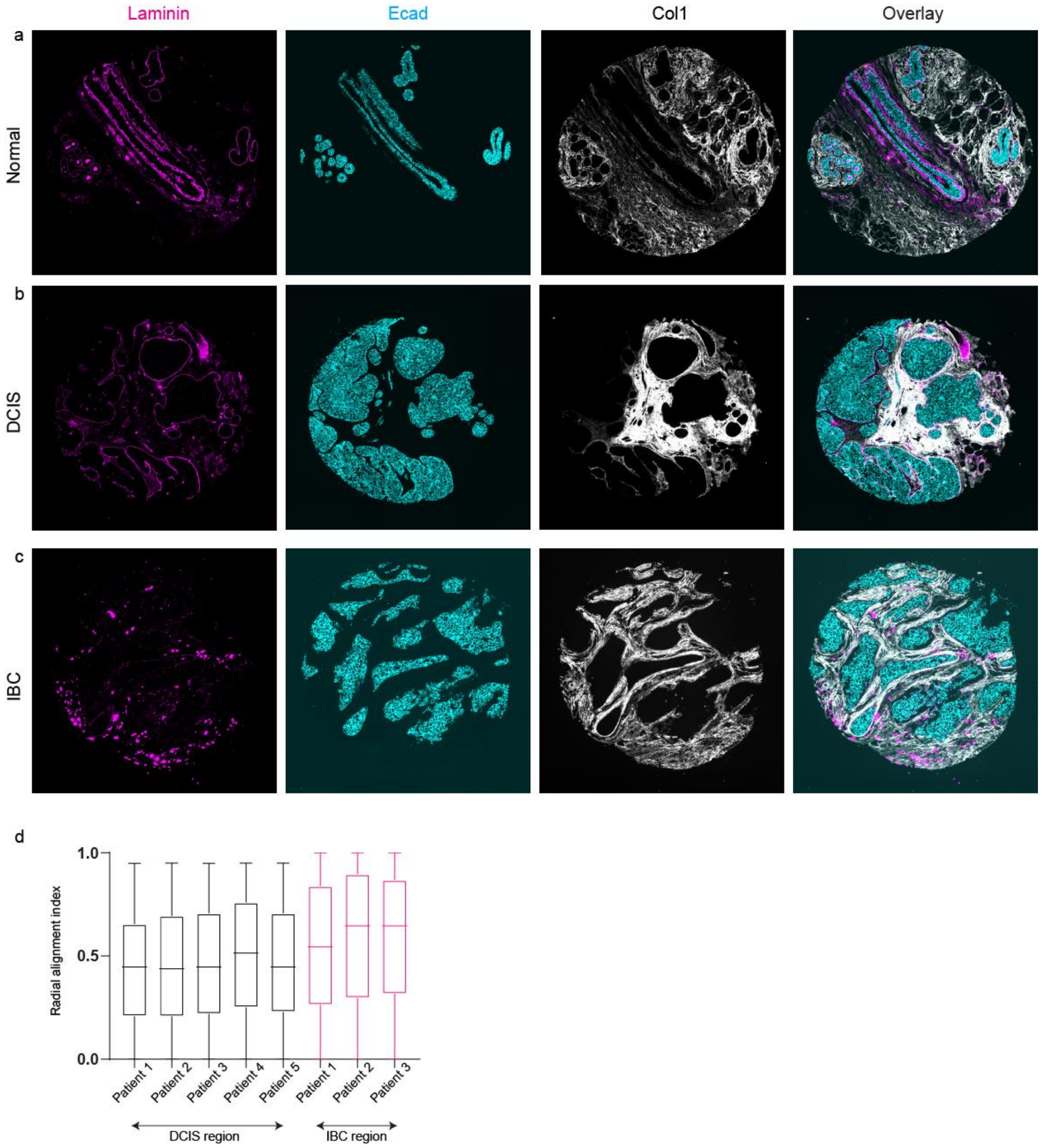


**Extended Figure 6: Fluorescent imaging of human histology sections and radial alignment index. a-c,** Fluorescent images of laminin, Ecad and col1 and the overlay of the fluorescent images in normal, DCIS and IBC regions of a human tumor. **b,** Radial alignment index around Ecad+ tumor cell clusters in DCIS and IBC regions of different patients.

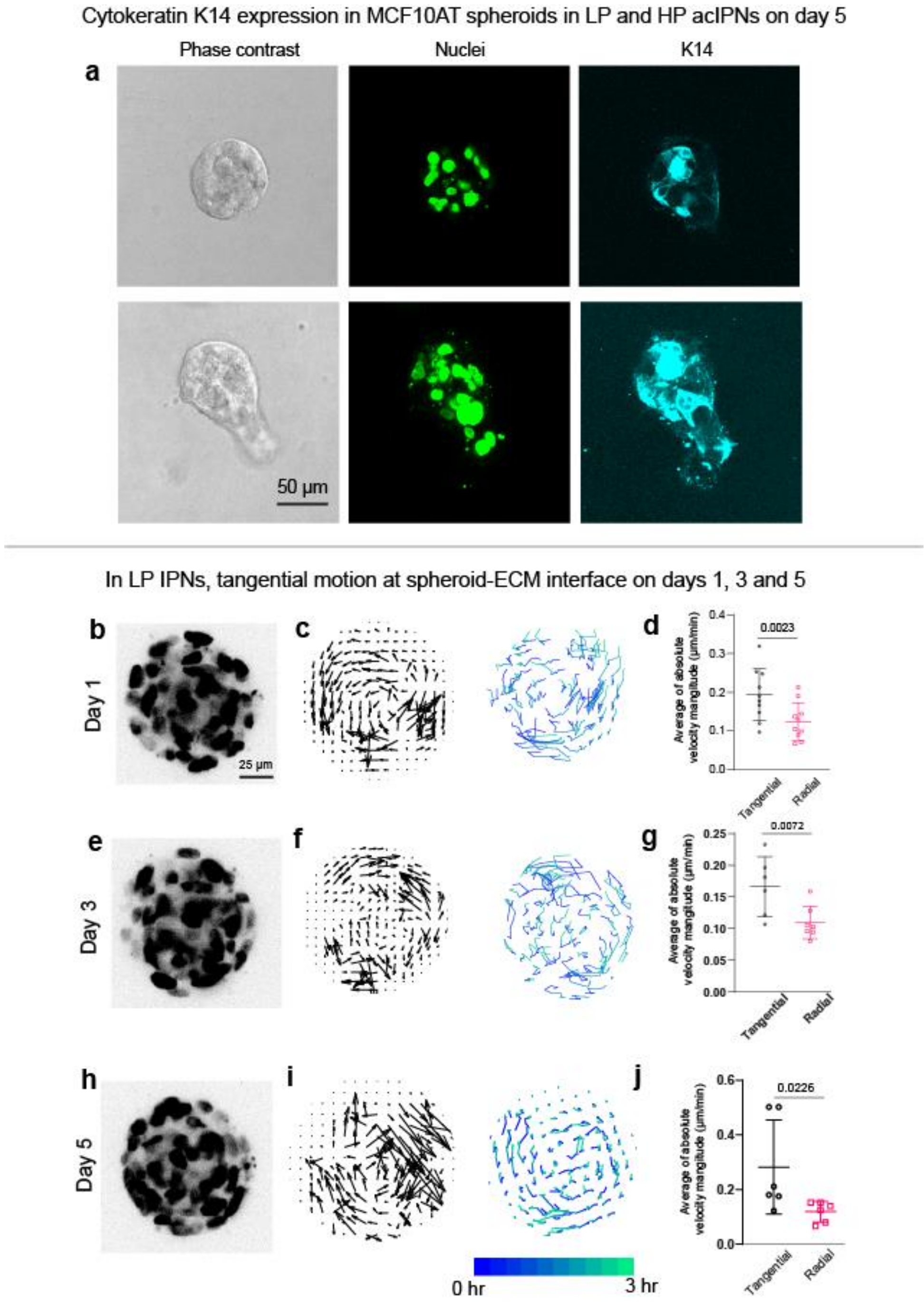


**Extended Figure 7: K14 expression, and swirling motion in LP acIPNs and in an orthotopic sarcoma xenograft model. a,** Phase contrast of an invading MCF10AT spheroids and scanning confocal images of H2B-GFP and K14 expression in the invading spheroid on day 5 in HP acIPN. **b**, Scanning confocal image of H2B-GFP nuclei in the equatorial plane of MCF10AT spheroids day 1 in LP acIPN. **c,** Instantaneous cell velocity field and cell trajectories over 3 hrs of time lapse imaging in the equatorial plane of MCF10AT spheroids on day 1 in LP acIPN. **c,** Average of absolute radial and tangential velocities near spheroid-ECM interface in the spheroid's equatorial plane on day 1 in LP acIPN (n=9; N=2). **e**, Scanning confocal image of H2B-GFP nuclei in the equatorial plane of MCF10AT spheroids day 3 in LP acIPN. **f,** Instantaneous cell velocity field and cell trajectories over 3 hrs of time lapse imaging in the equatorial plane of MCF10AT spheroids on day 3 in LP acIPN. **g,** Average of absolute radial and tangential velocities near spheroid-ECM interface in the spheroid's equatorial plane on day 3 in LP acIPN (n=6; N=2). **h**, Scanning confocal image of H2B-GFP nuclei in the equatorial plane of MCF10AT spheroids day 5 in LP acIPN. **i,** Instantaneous cell velocity field and cell trajectories over 3 hrs of time lapse imaging in the equatorial plane of MCF10AT spheroids on day 5 in LP acIPN. **j,** Average of absolute radial and tangential velocities near spheroid-ECM interface in the spheroid's equatorial plane on day 5 in LP acIPN (n=6; N=2). Panels **d, g, j -** Two-sided t-test.

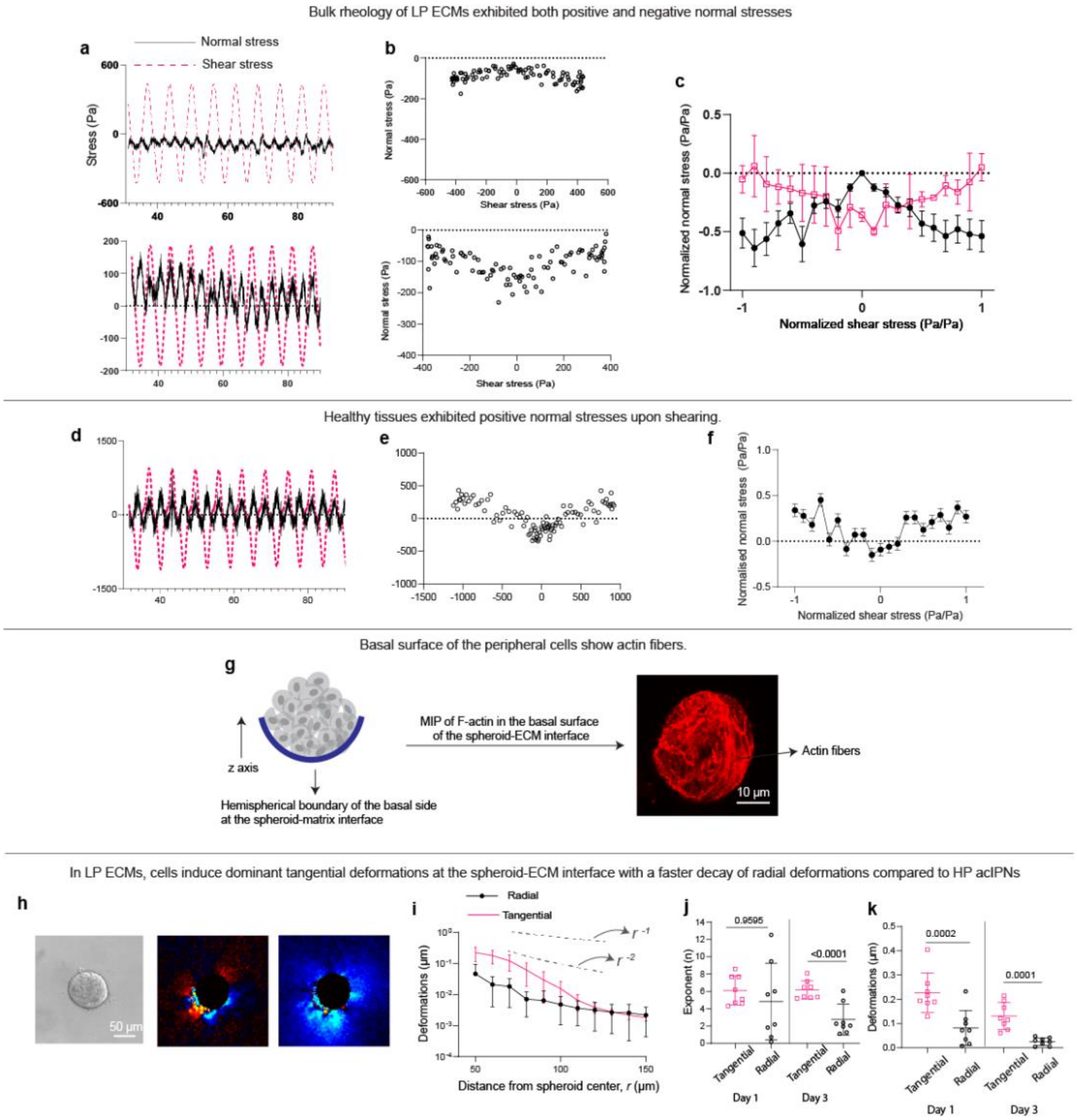


**Extended Figure 8: Bulk rheology, actin stress fibers, and ECM mechanical deformations in LP acIPNs. a-b,** Temporal plot of normal stresses and shear stresses during an oscillatory shear rheology of LP acIPNs at 20% strain amplitude, and the corresponding scatter plot between the normal stress and shear stress. **c,** Average of normalized normal stresses with respect to normalized shear stress (n=2 hydrogels for positive and negative normal stresses). **d-e,** Temporal plot of normal stresses and shear stress with time during an oscillatory shear rheology of a healthy breast tissue at 20% strain amplitude, and the corresponding scatter plot between normal and shear stresses. **f,** Average of normalized normal stresses with respect to normalized shear stress (n = 2 patients). **g,** (Left) Schematic depicting actin intensity projection of basal side of cells in the lower half of a cancer spheroid. (Right) Confocal images of actin at the basal side of the lower hemispherical part of a MCF10AT spheroid. **h,** Phase contrast of MCF10AT spheroid in LP acIPN, and heatmaps of radial and tangential deformations in LP acIPN on day 1. **i,** Line plots showing the average of radial and tangential deformations across different spheroids with respect to distance from the spheroid center on day 1. **j,** Exponents from power law fit of the line plots of radial and tangential deformations for days 1 and 3 in LP acIPNs. **k,** Average of radial and tangential deformations near spheroid-ECM interface in the equatorial plane of MCF10AT spheroids in LP acIPNs on days 1 and 3 (n=8; N=2). Panels **j**, **k** Two-sided t-test.

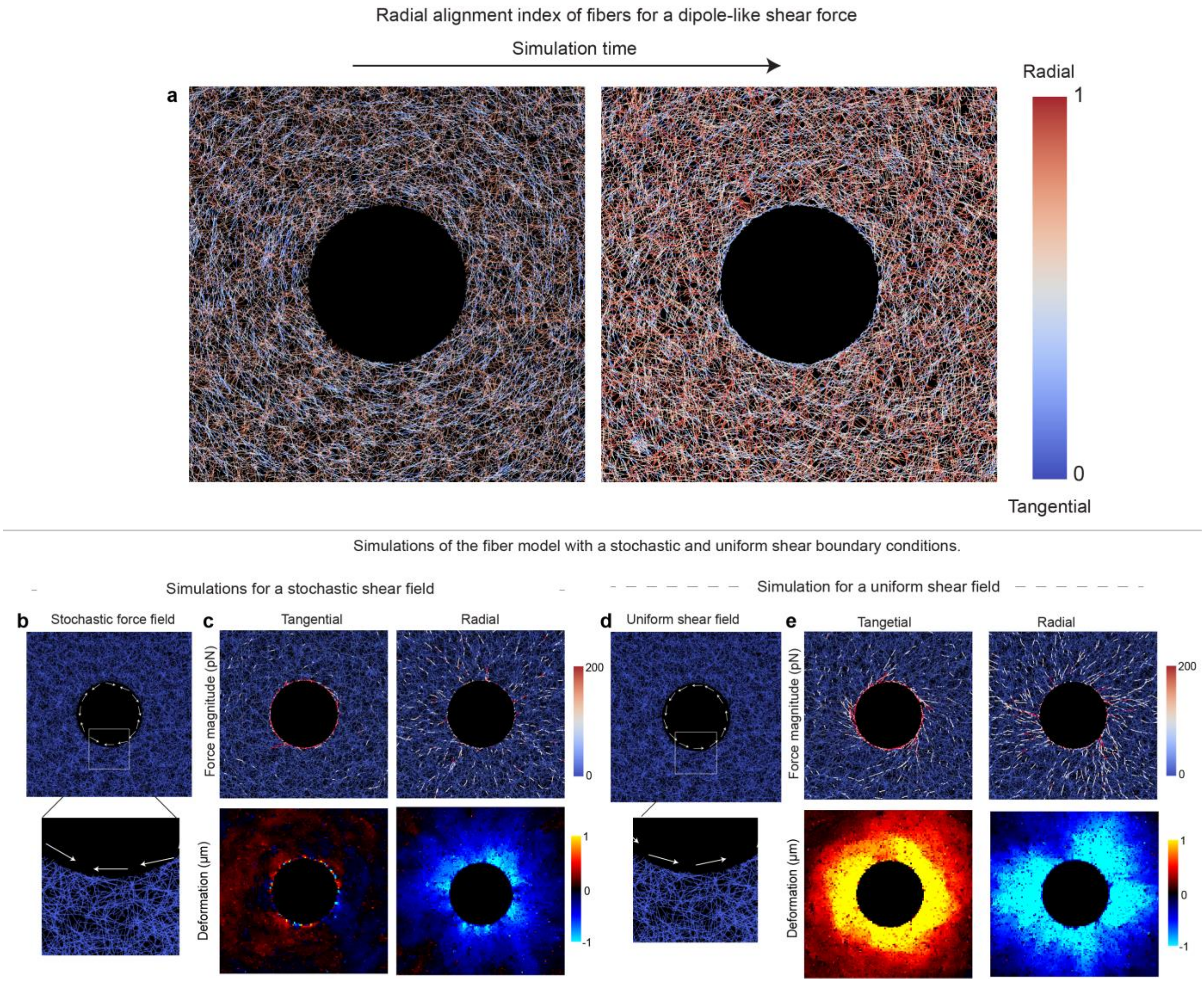


**Extended Figure 9: Fiber model simulations. a**, Snapshots of the radial alignment index at initial (left) and final (right) time points of the fiber simulation for a dipole-like shear force. **b**, Snapshot of boundary conditions for a stochastic shear field. White arrows indicate the direction of the shear force. **c**, Snapshots of force magnitude and deformations in tangential and radial direction for a stochastic shear field. **d**, Snapshot of boundary conditions for a uniform counterclockwise shear field. **e**, Snapshots of force magnitude and deformations in tangential and radial direction for a uniform shear field.

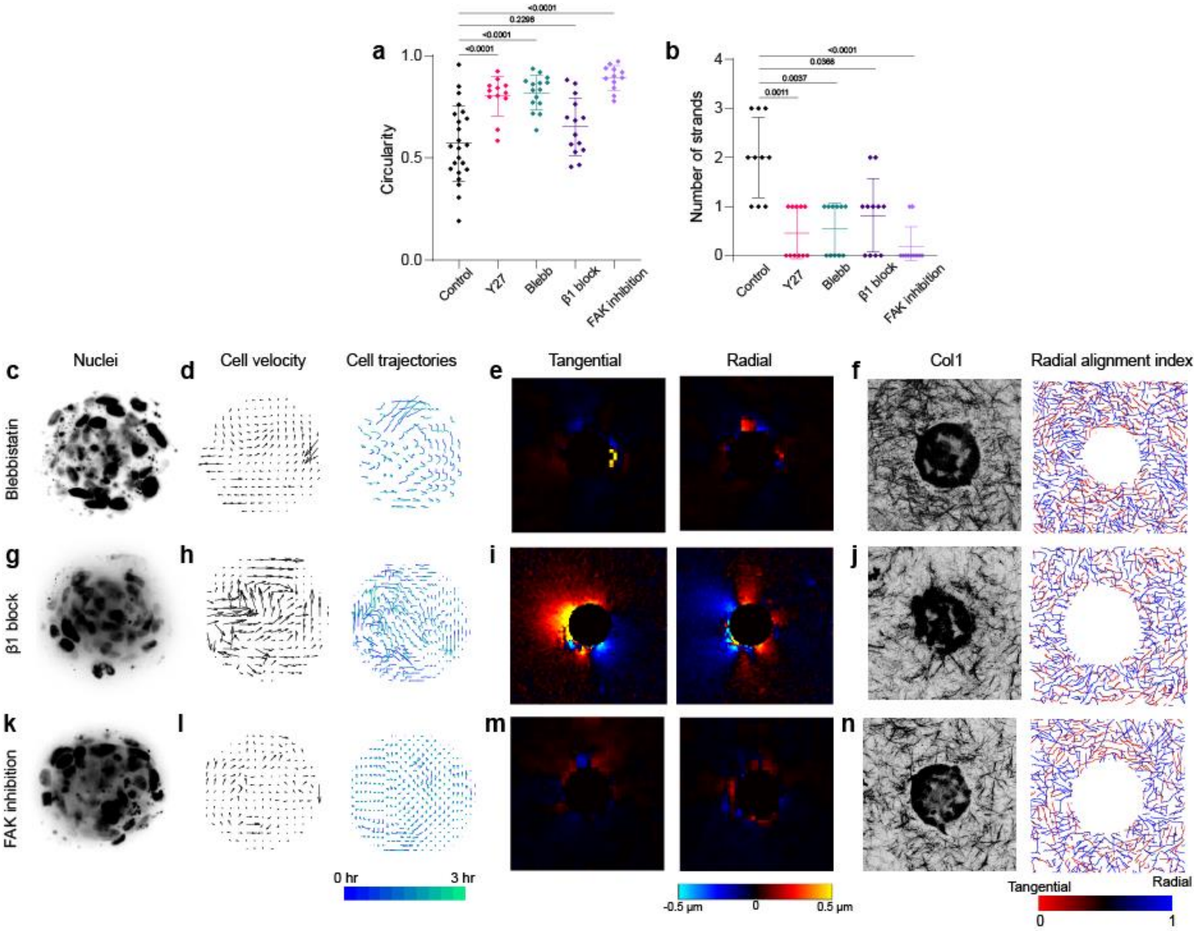


**Extended Figure 10: Role of cell contractility and cell-matrix adhesions on swirling motion and col1 alignment. a-b,** Circularity and number of collectively invading strands on day 5 in different conditions (n>7; N=3). **c,** Scanning confocal images of H2B-GFP nuclei in the equatorial plane of blebbistatin-treated MCF10AT spheroid on day 3 in HP acIPN. **d,** Instantaneous cell velocity field and cell trajectories over 3 hrs of time lapse imaging in the equatorial plane of blebbistatin-treated MCF10AT spheroids on day 3 in HP acIPN. **e,** Heatmaps of radial and tangential deformations in the equatorial plane of blebbistatin-treated MCF10AT spheroid on day 3. **f,** Scanning confocal reflectance of col1 around blebbistatin-treated MCF10AT spheroid on day 5 in HP acIPN and the corresponding heatmap of radial alignment index. **g,** Scanning confocal images of H2B-GFP nuclei in the equatorial plane of β1-blocking MCF10AT spheroid on day 3 in HP acIPN. **h,** Instantaneous cell velocity field and cell trajectories over 3 hrs of time lapse imaging in the equatorial plane of β1-blocking MCF10AT spheroid on day 3 in HP acIPN. **i,** Heatmaps of radial and tangential deformations in the equatorial plane of β1-blocking MCF10AT spheroid on day 3 in HP acIPN. **j,** Scanning confocal reflectance of col1 around β1-blocking MCF10AT spheroid on day 5 in HP acIPN and the corresponding heatmap of radial alignment index. **k,** Scanning confocal images of H2B-GFP nuclei in the equatorial plane of FAK-inhibited MCF10AT spheroid on day 3 in HP acIPN. **l,** Instantaneous cell velocity field and cell trajectories over 3 hrs of time lapse imaging in the equatorial plane of FAK-inhibited MCF10AT spheroid on day 3 in HP acIPN. **m,** Heatmaps of radial and tangential deformations in the equatorial plane of FAK-inhibited MCF10AT spheroid on day 3 in HP acIPN. n**,** Scanning confocal reflectance of col1 around FAK-inhibited MCF10AT spheroid on day 5 in HP acIPN and the corresponding heatmap of radial alignment index. Panels **a, b –** One-way ANOVA with Tukey's correction for multiple comparisons.
